# Comparative computational structural genomics highlights divergent evolution of fungal effectors

**DOI:** 10.1101/2022.05.02.490317

**Authors:** Kyungyong Seong, Ksenia V Krasileva

## Abstract

Elucidating the evolution of pathogen effector molecules is critical to understand infection mechanisms of fungal phytopathogens and ensure crop security. However, rapid diversification that diminishes sequence similarities between homologous effectors has largely concealed the roots of effector evolution. We predicted the folds of 26,653 secreted proteins from 21 species with AlphaFold and performed structure-guided comparative analyses on two aspects of effector evolution: uniquely expanded effector families and common folds present across the fungal species. Extreme expansion of nearly lineage-specific effector families was found only in several obligate biotrophs, *Blumeria graminis* and *Puccinia graminis*. The highly expanded effector families were the source of conserved sequence motifs, such as the Y/F/WxC motif. We identified additional lineage-specific effector families that include known (a)virulence factors, such as AvrSr35, AvrSr50 and Tin2. These families represented new classes of sequence-unrelated structurally similar effectors. Structural comparisons revealed that the expanded structural folds further diversify through domain duplications and fusion with disordered stretches. Sub-and neo-functionalized structurally similar effectors reconverge on regulation, expanding the functional pools of effectors in the pathogen infection cycle. We also found evidence that many effector families could have originated from ancestral folds that have been conserved across fungi. Collectively, our study highlights diverse mechanisms of effector evolution and supports divergent evolution of effectors that leads to emergence of pathogen virulence from ancestral proteins.

## Introduction

Fungal phytopathogens rely on secreted proteins termed effectors to suppress plant immunity, modify host cellular activities and successfully colonize the hosts (Lo Presti et al., 2015). This makes it critical to elucidate the molecular functions and biological roles of effectors for understanding infection mechanisms and ultimately protecting crops from the pathogens. However, effectors typically lack sequence similarity, functional annotations, and commonly shared sequence features (Sperschneider et al., 2015a, 2015b). This absence of sequence-based evolutionary guidance has hindered rationally selecting effector targets and illuminating effector functions, while genome sequencing of new phytopathogenic species has rapidly expanded the list of putative effectors.

Many effectors are unrelated by their primary sequences but share similar structures (de Guillen et al., 2015; Franceschetti et al., 2017; Saur et al., 2019; Spanu, 2017; Lazar et al., 2020; Seong and Krasileva, 2021; Yu et al., 2021; Outram et al., 2022). These effectors are major players in the battlefield of plant immunity and fungal pathogens. Plant intracellular immune receptors can evolve specificity towards these effectors to acquire resistance against the pathogens, as represented with MAX effectors (*Magnaporthe oryzae* Avrs and ToxB) (Ortiz et al., 2017; De la Concepcion et al., 2018; Guo et al., 2018; Białas et al., 2021). In turn, pathogens repeatedly lose and gain effectors to evade immune recognition (Yoshida et al., 2016; Kim et al., 2019; Latorre et al., 2020). Repeated appearance of sequence-unrelated structurally similar effectors and their contributions to pathogen virulence signify their importance. However, only a few classes of such fungal effector families have been discovered to date.

Divergent evolution was suggested to drive effector evolution (de Guillen et al., 2015). That is, a group of sequence-unrelated structurally similar effectors originated from a common ancestor but have lost detectable sequence similarity through rapid diversification. We proposed computational structural genomics as a framework to reveal such evolutionary connections obscured by sequence dissimilarity with predicted structures (Seong and Krasileva, 2021). The success of the methodology was exemplified by the identification of the MAX effector cluster, which could not be revealed by remote homology searches alone (Jones et al., 2021). With the availability of AlphaFold (Senior et al., 2020; Jumper et al., 2021), secretome-wide structure prediction and analysis has provided further insights. For instance, many important effectors from *Fusarium oxysporum* f. sp. *lycopersici* could be grouped into a few structural families, including Fol dual-domain (FOLD) effector family (Yu et al., 2021). In *Venturia inaequalis*, MAX effectors represent one of the most expanded families (Rocafort et al., 2022). Such structural analyses have reinforced the divergent evolution hypothesis in that pathogen virulence factors may have evolved through frequent duplications and functional diversification of ancient folds.

We proposed that computational structural genomics at a comparative scale would reveal novelty and commonality of effector and better elucidate effector evolution across diverse species in the fungal kingdom (Seong and Krasileva, 2021). To elucidate effector evolution at the structural level, we predicted with AlphaFold the folds of 26,653 secreted proteins from 14 agriculturally important fungal phytopathogens (Dean et al., 2012), 6 non-pathogenic fungi, and oomycete *Phytophthora infestans* as an outgroup. We focus on two aspects of effector evolution: uniquely expanded effector families and common folds present across the fungal species. We highlight how structural information overlaid on sequence-dissimilar effectors can provide insights into effector evolution.

## Results

### Structure prediction for fungal secretomes with AlphaFold

To perform comparative computational structural genomics, we predicted with AlphaFold the structures of 26,653 proteins collected from 21 species’ secretomes (Fig. 1; Table S1). This list of species includes agriculturally important phytopathogens with various lifestyles and host ranges that span across two phyla, Ascomycota and Basidiomycota (Dean et al., 2012). We added putatively saprotrophic, non-phytopathogenic species per order or subphylum as controls, and the oomycete *P. infestans* as an outgroup for its importance. We used pTM scores provided by AlphaFold as a global measure of the estimated precision of the predicted structures. The pTM score > 0.5 was used as a threshold to select reliably predicted folds as in our previous study (Seong and Krasileva, 2021). Some AlphaFold models only produce per-residue confidence scores (pLDDT), which range from 0 to 100, as a local accuracy measure, and the average pLDDT is used to evaluate the overall prediction quality. Although the average pLDDT scores correlate well with pTM scores (Fig. S1), the distribution of the average pLDDT scores tended to be skewed towards higher scores than that of the pTM scores (Fig. S2). Given the relationships between pLDDT and pTM, we found that pTM > 0.5 would roughly correspond to pLDDT > 70 (Fig. S3).

**Figure 1.**
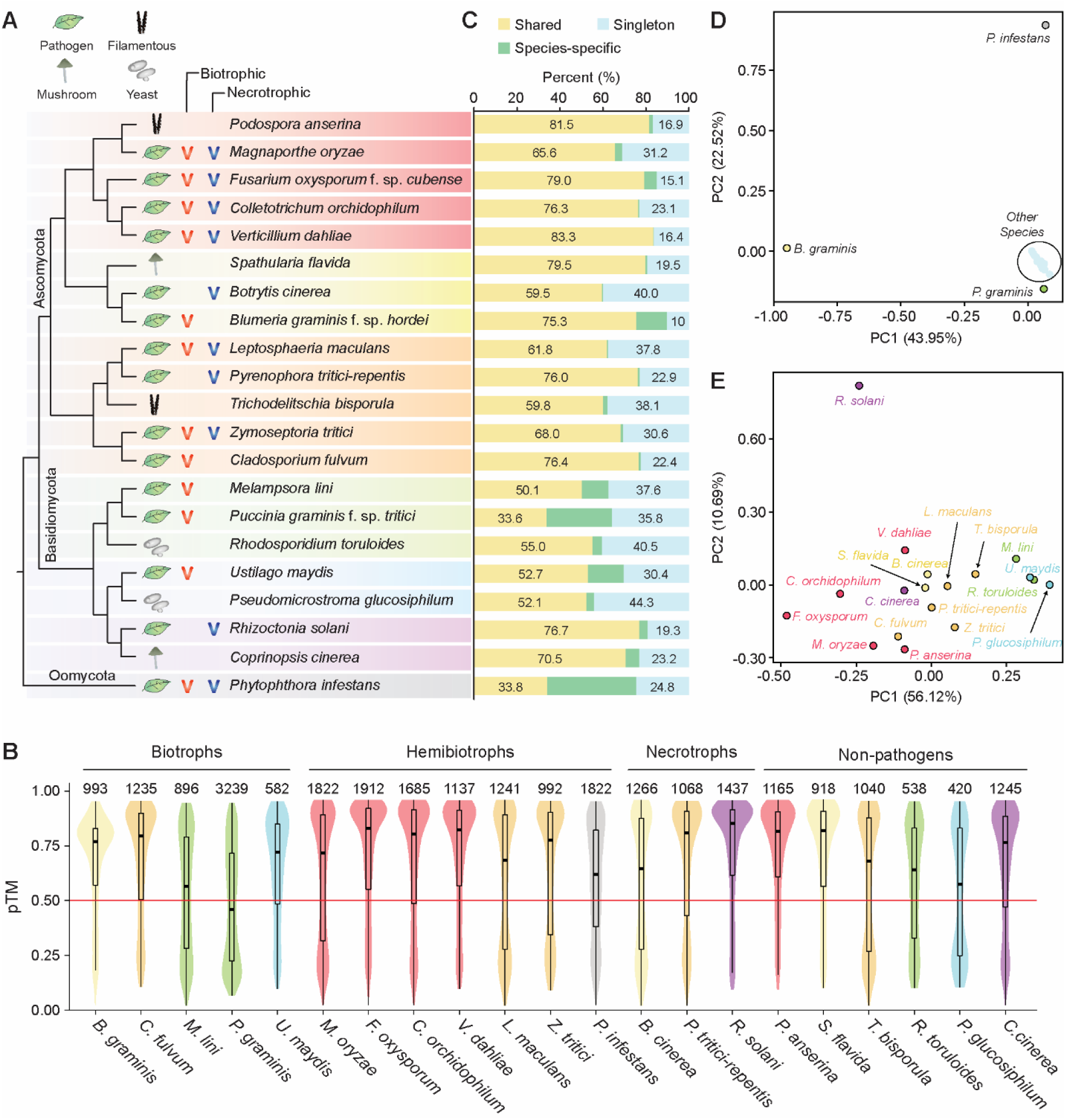
The design of comparative genomics study and statistics of structure prediction and secretome clustering. **A**. The hypothetical species tree reconstructed based on MycoCosm (Grigoriev et al., 2014) and lifestyles of the 21 species included in this study. The classification of fungal species (mushroom, filamentous and yeast) depended on their morphology. **B**. The distribution of pTM scores used to measure the quality of structure prediction. The total number of secreted proteins is indicated for each species. The species are grouped based on their lifestyles and ordered alphabetically. **C**. The proportion of proteins in shared, species-specific clusters or as singletons in whole-secretome clustering. The secretomes of the 21 species were clustered based on sequence and structural similarities. The clusters were categorized as ‘shared’ if the cluster members were found in more than one species, ‘species-specific’ if the cluster members originated from a single species, and ‘singleton’ if the protein does not belong to a cluster (Table S2). **D** and **E**. The principal component analysis (PCA) on the copy number variations of the clusters in the final clustering output. Singletons were not used for the analysis. The species that belong to the same class or subphylum are indicated with the same color, and the colors correspond to the background highlights given in A. **D**. All 21 species in this study were included. **E**. The three outliers, *Blumeria graminis, Puccinia graminis* and *Phytophthora infestans*, were removed, and the PCA was re-performed.

In comparison to the structural models produced in our previous study with TrRosetta for hemibiotrophic *M. oryzae* (Seong and Krasileva, 2021), the estimated precision of AlphaFold models were typically greater (Fig. S1; Fig. S4). However, even with the enhanced prediction performance of AlphaFold, only 55 additional protein structures were modeled by AlphaFold (Fig. S4). Moreover, 612 (33.5%) of *M. oryzae*’s secreted proteins missed by TrRosetta could also not be predicted by AlphaFold (Fig. S5). Overall, AlphaFold predicted from 47.0% (1,523 proteins in *Puccinia graminis* f. sp. *tritici*) to 81.5% (949 proteins in *Podospora anserina*) of the secreted proteins across the species in this study (Fig. 1B; Table S1). In total, 17,944 (67%) out of 26,653 proteins were modeled with the pTM scores > 0.5. The lifestyle of the species was not a determining factor of AlphaFold’s performance. For instance, AlphaFold performed well on the secretome of biotrophs, *Blumeria graminis* f. sp. *hordei* (*Bgh*), *Cladosporium fulvum* and *Ustilago maydis*, predicting about 75% of the secreted proteins (Fig. 1B; Table S1). Conversely, *P. graminis* f. sp. *tritici* (*Pgt*) and *Melampsora lini* were challenging targets. Relatively lower performance of AlphaFold was also observed in species with other lifestyles. This suggested that the varying performance of AlphaFold may be attributed to lineage-specific evolution that results in differing secretome inventory.

### Secretome clustering with sensitive sequence similarity searches and structural comparisons

To reveal evolutionary connections between secreted proteins, we clustered the secretome of individual species based on sequence and structural similarities. The similarity searches were performed sequentially for sequence-to-sequence with BLASTP, sequence-to-profile with HHblits, profile-to-profile with HHsearch, and structure-to-structure with TM-align as in our previous study (Seong and Krasileva, 2021). The proportion of clustered proteins ranged from 29.0% in *Pseudomicrostroma glucosiphilum* to 76.2% in *Bgh* (Table S2 and S3). For a comparative analysis, we clustered the entire secretomes of the 21 species used in this study (Table S2 and S4). Overall, 7,207 (27%) proteins, the majority of which do not have predicted structures, still remained as singletons (Fig. S6; Table S2). However, 4,087 (15%) proteins initially found as singletons in the species-wide clustering had sequence or structure-related proteins in other species and could be assigned to the clusters (Fig. S7). In total, 19,446 (73%) of the secreted proteins had at least one homolog or analog within or outside the species’ secretome. We classified the proteins into ‘shared’ if they belong to clusters of two or more species and ‘species-specific’ otherwise (Fig. 1C). Except for the outgroup, *P. infestans*, only *Bgh, M. lini, Pgt* and *U. maydis* displayed a relatively high proportion of species-specific secreted proteins (>10%) (Table S2). Most proteins belonged to the shared clusters, possibly suggesting that many proteins may have common ancestral origins.

### The composition of fungal secretomes largely reflect evolutionary relationships between the species

We then examined the copy number variations within the clusters with principal component analysis (PCA) to examine any patterns associated with the clusters (Table S5). The clusters generated only based on simple sequence similarity search with BLASTP revealed the species clustered largely based on their evolutionary distances, except for the two outliers, *Bgh* and *Pgt* (Fig. S8). However, the first two principal components (PCs) could only explain 37.5% of the variance. On the other hand, with the final clusters guided by sensitive sequence similarity searches and structural comparisons, the first two PCs captured about 66.8% variance (Fig. 1E and 1F). The structure of the two fungal outliers and other species generally spread based on the evolutionary distances remained. This suggests that except for some obligate biotrophs that may undergo distinct evolution, evolutionarily closely related species would likely have more similar compositions of secretomes, but even such evolutionary connection may be masked by sequence dissimilarities between homologs.

### Nearly or entirely species-specific effector families exist only in some phytopathogens

We examined nearly (>80%) or entirely species-specific clusters with known virulence and avirulence factors collected from the literature and PHI-base (Table S5 and S6) (Urban et al., 2022). Consistent with the observation that *Bgh, Pgt* and *P. infestans* were the outliers, only these species had nearly or entirely species-specific, highly expanded effector families with 100 or more members (Fig. 2A). Many clusters, such as Cluster 29 (119 members), 31 (118 members), 40 (87 members) and 62 (58 members), were also nearly exclusive to *Bgh* or *Pgt*; however, no virulence factors related to these clusters have been yet studied to our knowledge (Fig. S9). The Tin2-like effector family in *U. maydis*, as well as MAX effectors and ADP-ribosyl transferase (ARTs) families in *M. oryzae* were followed but only had about 30 members (Fig. 2A). In other fungal phytopathogens, we did not observe any nearly or entirely species-specific effector families with comparable sizes (>15 members). This result highlighted unique evolution of fungal obligate biotrophs, *Bgh* and *Pgt*, with extreme expansions of a few effector families.

**Figure 2.**
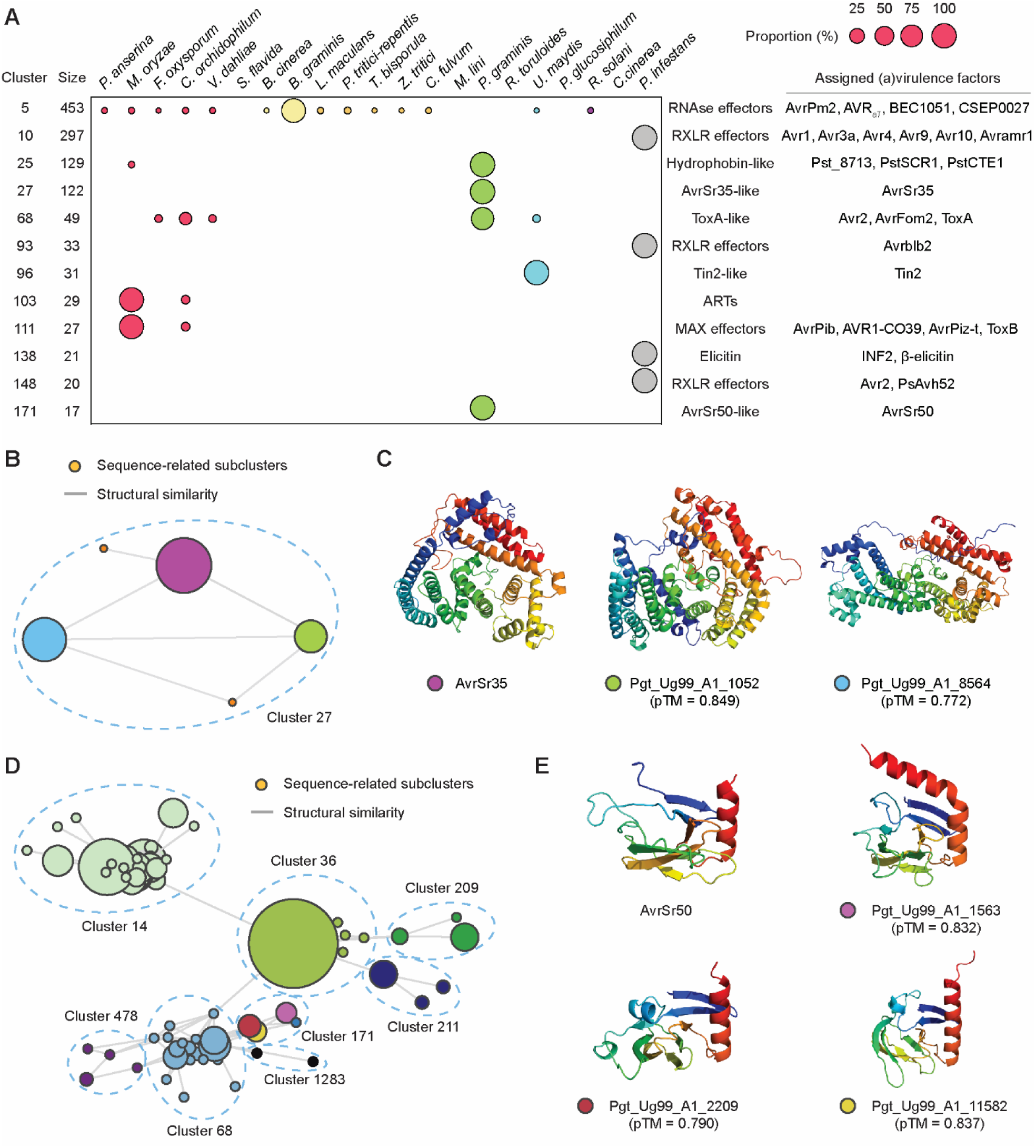
The expanded effector families in phytopathogens and new classes of sequence-unrelated structurally similar effectors. **A**. The nearly or entirely species-specific effector families with putative functions or known (a)virulence factors. The relative compositions of individual species in each cluster are indicated with circles of varying sizes. Only a subset of known (a)virulence factors are indicated (Table S5). **B** and **D**. The network of Cluster 27, as well as Cluster 171 and other related clusters. Each node represents a sequence-related subcluster or singleton, and the edges indicate structural similarity between the subclusters or singletons. The size of the nodes varies, depending on the number of subcluster members. **C** and **E**. The experimentally determined or predicted structures from *Puccinia graminis* selected from the subclusters in Cluster 27 and 171. The colored dots in the labels indicate the membership of the sequences. These colors correspond to those of nodes in B and D.

### Known virulence and avirulence factors may represent novel classes of sequence-unrelated structurally similar effectors

We next examined sequence and structural similarity between the members and the virulence factors assigned to the clusters. We found a common trend that the similarity between the cluster members and virulence factors can be detected only at the structural level. For instance, Cluster 27, to which AvrSr35 belongs, was composed of three sequence-related subclusters and two singletons connected by structural similarity (Fig. 2B and 2C). Cluster 171 specific to *Pgt* includes AvrSr50 (Fig. 2A). This cluster is interwound with other clusters in a complex manner, such as Cluster 68 with ToxA from *Pyrenophora tritici-repentis* and Avr2 from *F. oxysporum*, and Cluster 1283 with AvrL567 from *M. ilni* (Fig. 2D), that contain β-sandwich-like folds (Fig. S10) (Sarma et al., 2005; Wang et al., 2007; Di et al., 2017). However, the structural comparison between experimentally determined AvrSr50 (Ortiz et al., 2022) and members in Cluster 171 supported greater structural similarity between the sequence-unrelated secreted proteins (Fig. 2E). Together, the analysis suggested that sequence-unrelated structurally similar effectors have repeatedly evolved, and novel classes detected from structural similarity can include known avirulence and virulence factors.

### Some species-specific, highly expanded effector families may be the source of conserved high frequency motifs

The largest of nearly species-specific, highly expanded effector clusters was the RNAse-like effector family, composed of 453 members (Fig. 2A). Although other fungal species had a few RNAse-like proteins, 426 members were encoded in the *Bgh*’s genome, representing 43% of the *Bgh*’s secretome. As our clustering parameters were relatively stringent, we adopted the previously used parameters to recover the RNAse supercluster (Seong and Krasileva, 2021). We could retrieve 29 additional *Bgh*’s secreted proteins (4 clusters and 11 singletons) into the supercluster (Fig. 3A; Table S7). A previous study curated 491 candidates for secreted effector proteins (CSEPs) and grouped them into 72 gene families and 84 singletons based on sequence similarity (Pedersen et al., 2012). The authors reveal that 15 different families and 7 CSEP singletons were likely RNAse. We compared the membership of the CSEPs to the RNAse supercluster and found that 60 CSEP gene families and 41 CSEP singletons belong to the supercluster (Fig. 3A; Table S7). In other words, 70% of the CSEPs are putatively RNAses. Similarly, another previous study revealed that highly diverse putative effector groups in *B. graminis* share conserved Y/F/WxC motifs in the first 45 amino acids of the full-length proteins (Godfrey et al., 2010). We found that 371 (91.4%) out of the 406 Y/F/WxC motif-containing proteins belong to the RNAse-like effector supercluster (Fig. 3B; Table S7). For other secreted proteins than the putative RNAses, we did not observe any enrichment of Y/F/WxC motifs (Fig. S11).

**Figure 3.**
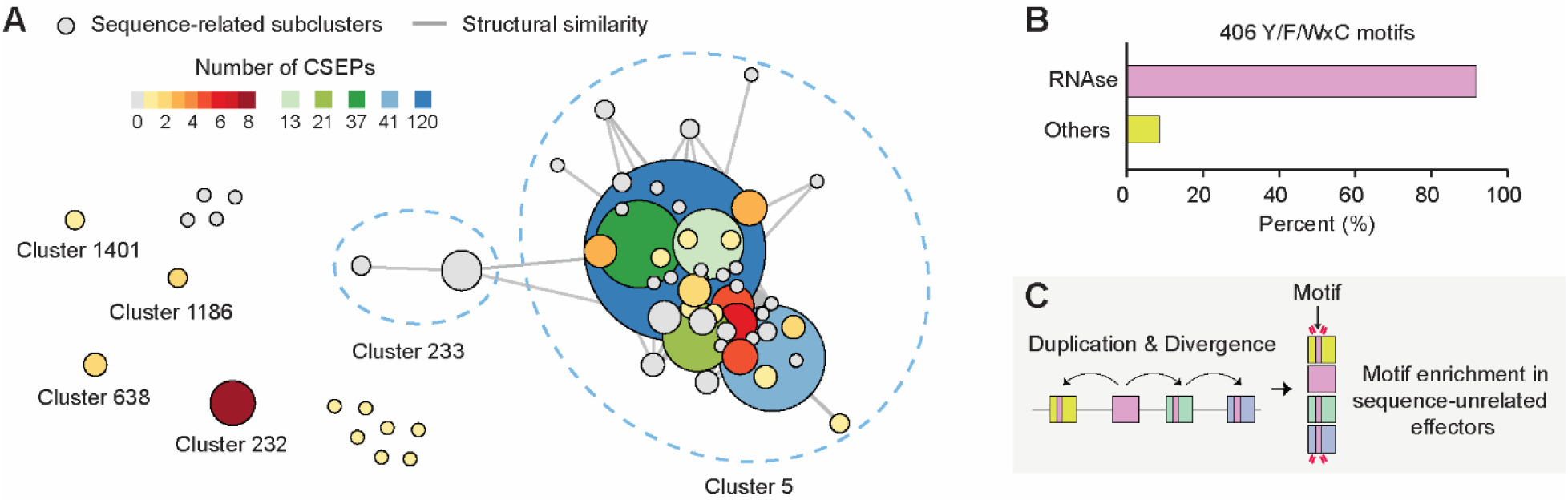
Highly expanded RNAse effector family and the emergence of Y/F/WxC-motifs in *Blumeria graminis*. **A**. The network of the RNAse supercluster. Each node represents a sequence-related subcluster or singleton, and the edges indicate structural similarity between the subclusters or singletons. The size of the nodes varies based on the number of members. The subcluster is colored to indicate the number of candidates for secreted effector proteins (CSEPs). Clusters or singletons, other than Cluster 5 and 233, were retrieved into the supercluster by lowering the stringency for clustering. **B**. The proportion of the Y/F/WxC motif-containing proteins in the RNAse supercluster or other clusters and singletons. **C**. The proposed explanation for the emergence of high-frequency conserved sequence motifs.

Similarly, we found that the Y/F/WxC motifs suggested to be present in *Pgt* were particularly enriched in Cluster 25 (Fig. S12) (Godfrey et al., 2010). In this sequence-unrelated structurally similar effector family, 90 (70%) out of 128 *Pgt* members contained the motif. Together, our data suggests that a high frequency sequence motif was unlikely to emerge by unrelated proteins independently and repeatedly acquiring the motif. Instead, divergent evolution of related proteins that diminishes sequence similarities more likely explains the presence of high frequency motifs only in some pathogenic species (Fig. 3C).

### Some effectors evolve from conserved fungal proteins to form a sequence-unrelated, structurally similar effector family

The largest, nearly species-specific effector family (Cluster 25) in *Pgt* lacked any functional annotations but consistently displayed structural similarity to hydrophobins (Figure 2A; Table S4). Investigating the structural similarity search results, we uncovered that Cluster 25 is related to Cluster 52 in which most fungal species used in this study had members and the majority of the cluster members were annotated as fungal hydrophobins at the sequence level (Fig. 4B). Nonetheless, the extreme sequence divergence that diminished sequence similarity between homologs was much more frequent in Cluster 25, as the majority of the nodes was connected only by structural similarity. Three virulence factors, PstCTE1, PstSCR1 and Pst_8713 from *Puccinia striiformis* f. sp. *tritici* show no detectable sequence similarity by BLAST; however, they displayed structural similarity and belonged to the hydrophobin-like cluster (Fig. 4C). Interestingly, while PstSCR1 was shown to be an apoplastic effector (Dagvadorj et al., 2017), PstCTE1 was suggested to localize in chloroplast (Andac et al., 2020) and Pst_8713 in cytoplasm and the nucleus (Zhao et al., 2018), potentially reflecting functional divergence. Together, our data suggest that sequence-unrelated structurally similar effector groups that may seem novel could have originated from conserved fungal proteins (Fig. 4A). Potentially, rapid sequence evolution for functional divergence and subsequent acquisition of new virulent functions may be driving the emergence of many sequence-related subclusters, the entire connectivity of which can be only discovered by structural comparisons.

**Figure 4.**
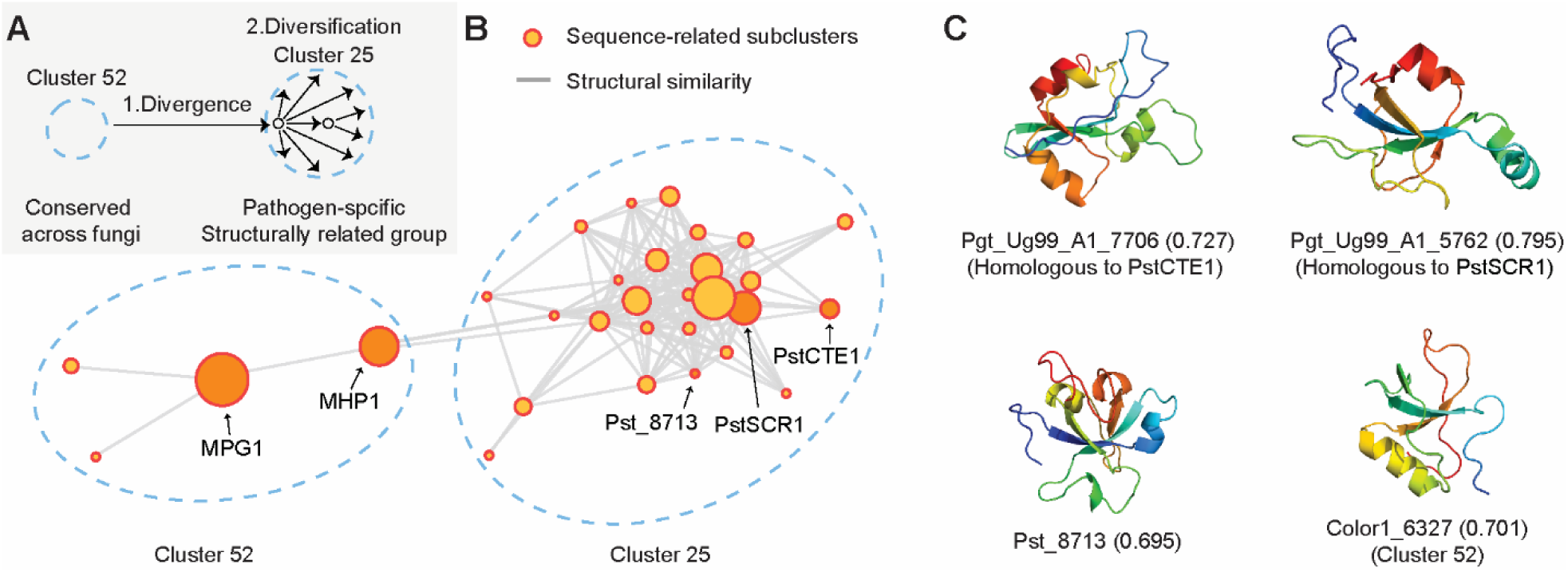
The emergence and expansion of hydrophobin-like effector family in *Puccinia graminis*. **A**. The proposed explanation for the emergence of the hydrophobin-like effector family in *Puccinia graminis*. **B**. The network graph of Cluster 25, nearly exclusive to *P. graminis*, and Cluster 52, present in most fungal species in this study. Each node represents a sequence-related subcluster or singleton, and the edges indicate structural similarity between the subclusters or singletons. The size of the nodes varies based on the number of members. The membership of the known virulence factors is indicated. **C**. Selected structures of the hydrophobin-like effector families. The top two structures are from *P. graminis*, while Pst_8713 and Color1_6327 are from *Puccinia striiformis* and *Colletotrichum orchidophilum*, respectively. In parentheses are pTM scores for the predicted structures.

### A genomic array of seemingly unrelated proteins could have resulted from rapid sequence evolution and subcluster expansion of related effectors

Previous studies analyzed a 40 kb genomic segment in chromosome 19 in *U. maydis*, that contains 23 secreted effectors comprising 5 gene families and multiple singletons (Kämper et al., 2006; Brefort et al., 2014). The deletion of this segment abolished the characteristic tumor formation of *U. maydis*, and Tin2 was identified as an important virulence factor that possibly alters the anthocyanin pathway in the plant hosts and reduces plant immune capabilities (Tanaka et al., 2014; Lanver et al., 2017). Our structure prediction and clustering suggest that the seemingly unrelated secreted proteins in the genomic segment, in fact, share structural similarity (Fig. 5A and 5B; Fig. S13). Furthermore, these Tin2-like effectors form the largest, species-specific sequence-unrelated structurally similar effector family in *U. maydis* (Fig. 2A). Brefort et al. reported that there were no paralogs on other chromosomes (Brefort et al., 2014). However, structural similarity searches revealed additional Tin2-like effectors duplicated and translocated into chromosomes 5 and 20 (Fig. 5B). A plausible explanation for such organizations of sequence-unrelated structurally similar effectors is frequent subcluster expansions after sequence divergence (Fig. 5C). That is, after a duplication of an ancestral Tin2-like effector occurs, one paralog rapidly diverges, losing sequence similarity. Subsequent tandem duplications may then expand subclusters composed of paralogs that maintain sequence similarity in proximity. This suggests that an array of seemingly unrelated proteins in pathogen genomes may have resulted from a single ancestral protein, and this can be only revealed at the structural level.

**Figure 5.**
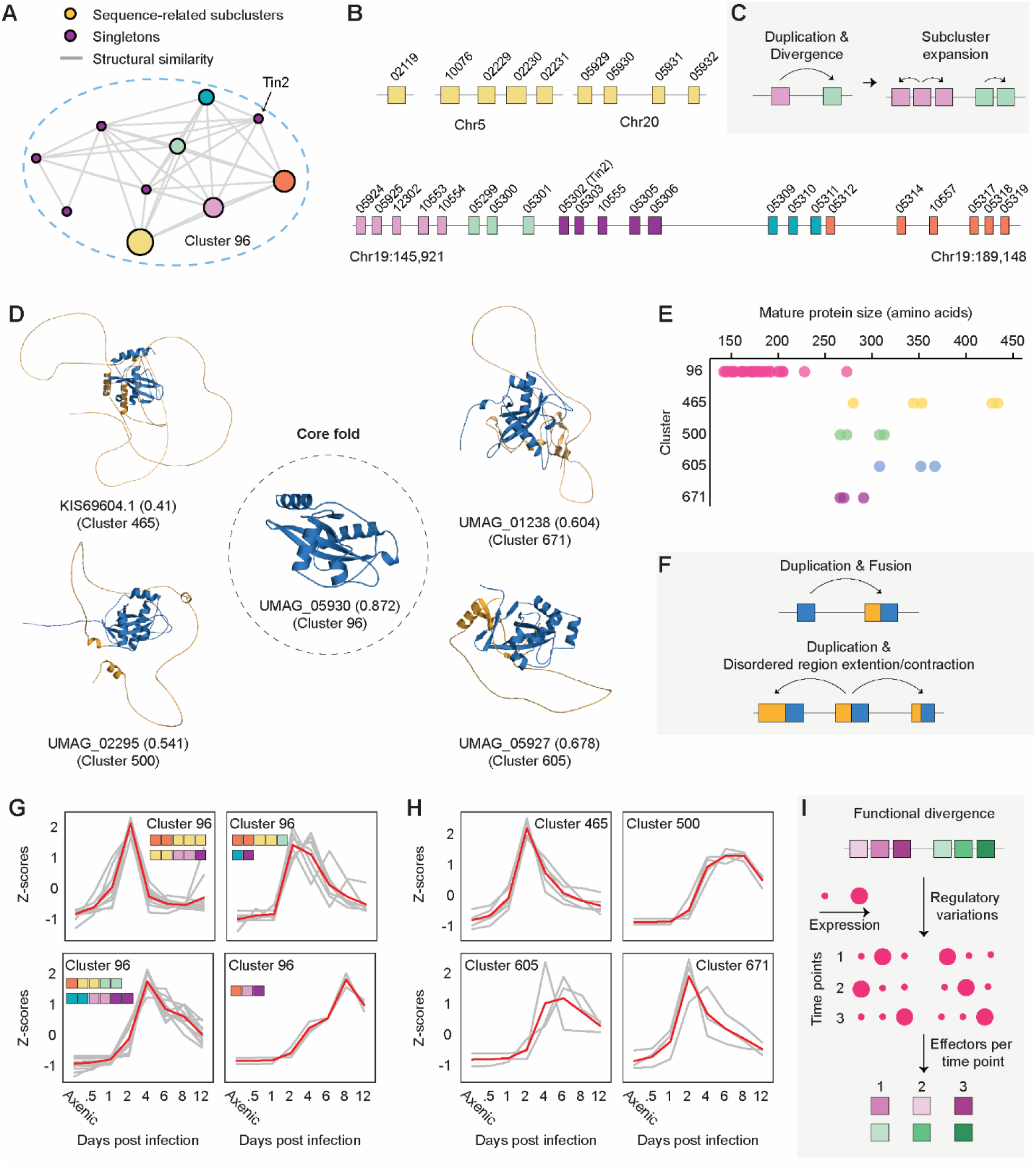
The evolution of Tin2-like effectors in *Ustilago maydis*. **A**. The network graph of Cluster 96, exclusive to *U. maydis*. Each node represents a sequence-related subcluster or singleton, and the edges indicate structural similarity between the subclusters or singletons. The size of the nodes varies based on the number of members. The membership of the known virulence factors is indicated. **B**. The genomic segments that include the members of Cluster 96. The colors, except purple, indicate the memberships given in A. The purple boxes represent singletons determined by sequence similarity searches. A few secreted proteins in this cluster are not depicted as they do not belong to Cluster 96. **C**. The proposed mechanism of the Tin2-like effector expansion. **D**. The selected structures from *U. maydis* that contain the Tin2 fold. The core Tin2 fold is colored in blue, and the disordered stretch in orange. The pTM scores are indicated in the parentheses, and the relatively lower pTM scores are attributed to the disordered stretches that do not adopt single rigid structures. **E**. The length distributions of the Tin2 fold-containing clusters. **F**. The proposed mechanism of the fusion between disordered regions and the Tin2 fold. **G**. The expression profile of Cluster 96 members. The members were grouped based on their similar expression patterns determined by hierarchical clustering. The membership of the sequences is indicated with colored boxes which correspond to the sequence-related subclusters given A and B. **H**. The expression profiles of Tin2 fold-containing disordered fusion proteins. **I**. The explanation for regulatory convergence of sequence-unrelated structurally similar effectors

### The core effectors diversify through fusing into disordered regions and extending and contracting them

Structural similarity searches of Tin2-like effectors indicated the presence of additional *U. maydis*-specific clusters in which the members may adopt similar folds. Upon visualizing the structures, we found that the members in these small clusters have the core Tin2 fold surrounded by long disordered stretches (Fig. 5D). In accordance, sequence-based disordered region prediction with IUPred2A (Mészáros et al., 2018) consistently supported that the N-terminal regions are intrinsically disordered with an abundance of glycines and prolines preventing secondary structure formation (Fig. S14). As the disordered regions could be misannotated, we relied on public transcriptomic data to confirm the gene models (Lanver et al., 2018). Even though the lengths of the mature proteins varied within the cluster and between the clusters (Fig. 5E), the single-exon gene models of the fusion proteins were supported either by de novo transcriptome assembly or transcriptome mapping (Fig. S15). Moreover, the expression of the fusion proteins was regulated throughout the infection cycle, and some of them displayed a high level of expression (Table S8), indicating that they would have functional roles. Together, this supported that the ancestral Tin2 fold was fused into a disordered region (Fig. 5F), and the extension and contraction of the disordered regions were followed after subsequent duplication events for possible diversification.

### Functionally diversified sequence-unrelated structurally similar effectors may converge on regulation

As the high-quality transcriptomic data is available for *U. maydis*, we examined the expression profiles of the Tin2-like effectors (Lanver et al., 2018). Hierarchical clustering of the core Tin2-like cluster (Cluster 96) revealed four distinct expression patterns with members from different sequence-related subclusters (Fig. 5B and 5G; Fig. S16). This suggests that the members in sequence-related subclusters underwent distinct regulatory mutations (Fig. 5I), and the functionally diversified Tin2-like effectors eventually re-converged in effector regulation, diversifying functional pools of effectors. On the contrary, the Tin2 fusion proteins, that have maintained sequence similarity among the members in the same cluster, tended to display similar expression profiles by clusters (Fig. 5H), possibly complementing the core Tin2-like effectors’ roles.

### Non-pathogenic species encode secreted proteins related to many virulence and avirulence factors

We next examined clusters that are not specific to phytopathogens (Fig. 6A; Table S5). Despite the presence of known avirulence and virulence factors, many clusters included members from non-phytopathogens, based on sensitive sequence similarity searches and structural comparisons. For instance, a large commonly shared cluster without a definitive role assigned to secreted proteins is Cluster 12 (Fig. 6B). Extracellular Ecp2 from *Cladosporium fulvum* with necrosis-inducing factor domain (PF14856) exists in this cluster. Nonetheless, about 30% of the members originated from the non-pathogenic species. The analysis of the network indicated that sequence-unrelated structural similarity was not necessarily a unique feature of phytopathogens, and identifying related proteins required structural comparisons for non-pathogenic species (Fig. 6C). Interestingly, in the sequence-related subcluster that only includes members from *Pgt* and *M. lini*, two yeast killer toxin-like domains were fused in a single protein (Fig. 6D and 6E). This fusion protein was supported by transcriptomic data (Chen et al., 2017) and appeared to be expanded in *Pgt* (Table S4). Collectively, such distinct evolution may reflect differing evolutionary pressures on this sequence-unrelated structurally similar groups from which virulence factors could evolve. That is, similarly to the RNAse-like and hydrophobin-like effector families in *Bgh* and *Pgt*, virulence factors may originate by divergent evolution of inherited secreted proteins, as an outcome of adaptation.

**Figure 6.**
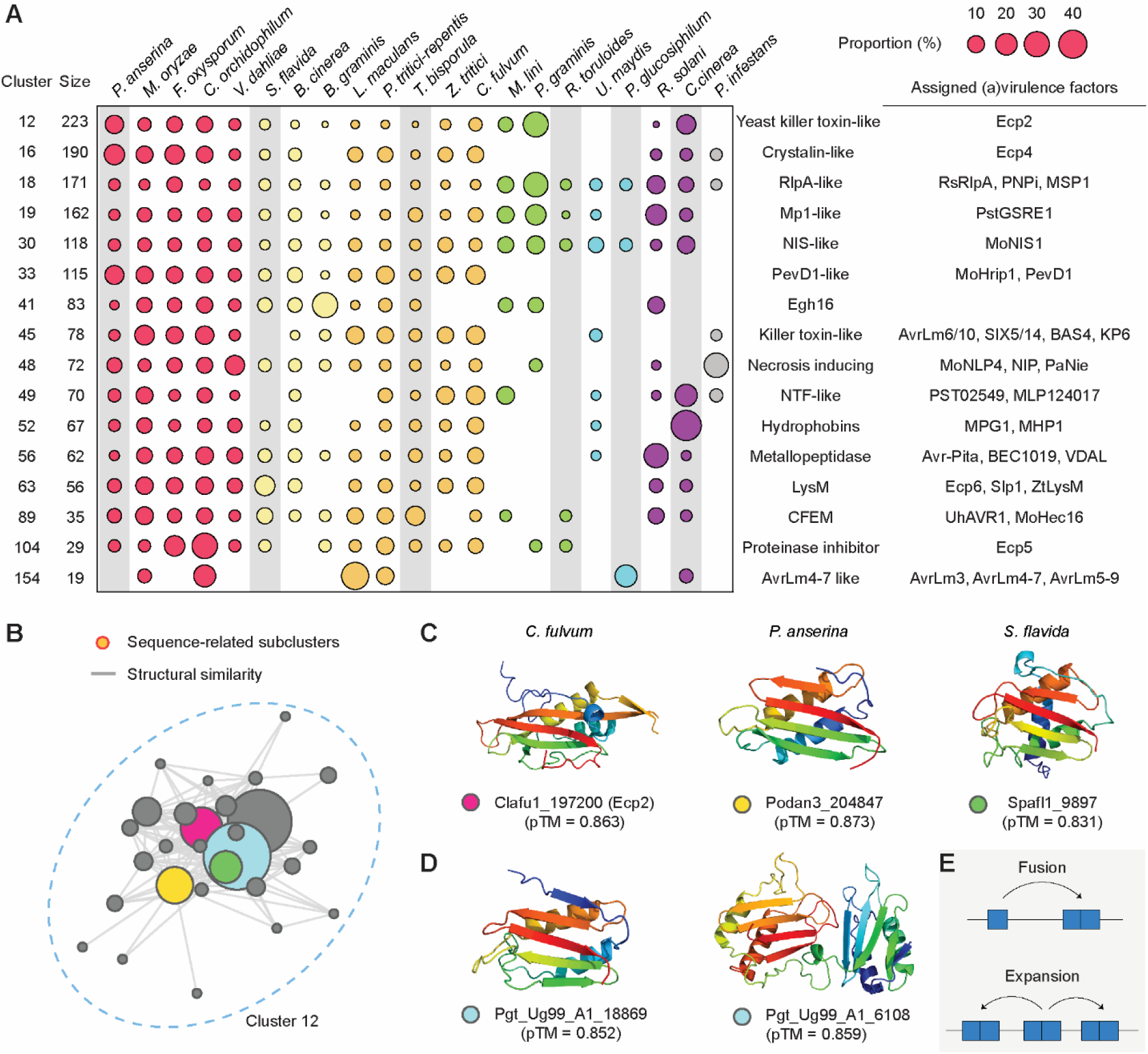
Divergent evolution of the commonly shared clusters. **A**. The putative effector families shared between phytopathogens and non-phytopathogens. The relative compositions of individual species in each cluster are indicated with circles of varying sizes. Only a subset of known (a)virulence factors are indicated (Table S5). Non-phytopathogenic species are highlighted with gray boxes. **B**. The network graph of Cluster 12. Each node represents a sequence-related subcluster or singleton, and the edges indicate structural similarity between the subclusters or singletons. The size of the nodes varies based on the number of members. The light blue subcluster contains secreted proteins only from *Puccinia graminis* and *Melampsora lini*. **C**. The selected predicted structures from different sequence-related subclusters. The membership of the secreted proteins is indicated with colored dots that correspond to the subclusters given in B. **D**. The selected predicted structures from *P. graminis*. These proteins belong to the light blue subcluster specific to *P. graminis* and *M lini*. **E**. The explanation for the emergence of novel dual-domain proteins in *P. graminis*.

## Discussion

Primary sequences of many fungal effectors cannot provide sufficient information about the effectors. Although pioneering tools, such as effectorP (Sperschneider and Dodds, 2022), guide the selection of effectors through the classification of secreted proteins based on the features of known effectors, they do not illuminate evolutionary or functional information. Computational structural genomics offers more intuitive information about the effectors through structural similarity to the existing and novel effector families. A comparative study further extends the evolutionary context to a broader scale and reveals additional information about effectors that the studies on single species may not capture. Through this study, we demonstrate the advantages of comparative computational structural genomics and how this method can reveal novel evolutionary insights about effectors masked by their sequence dissimilarities.

Our study primarily underscores divergent evolution of fungal effectors (Fig. 7). In this model, a protein shared in a common ancestor of non-phytopathogenic and phytopathogenic species can evolve to form sequence-unrelated structurally similar effector groups (Fig. 7A). After a duplication event of the ancestral protein, a paralog rapidly diverges in primary sequences and loses sequence similarity to the other paralog, while maintaining detectable structural similarity. Such processes occur repeatedly, eventually giving rise to multiple effector groups that are seemingly unrelated to each other. The connections of these groups can be only elucidated by the structural comparison.

**Figure 7.**
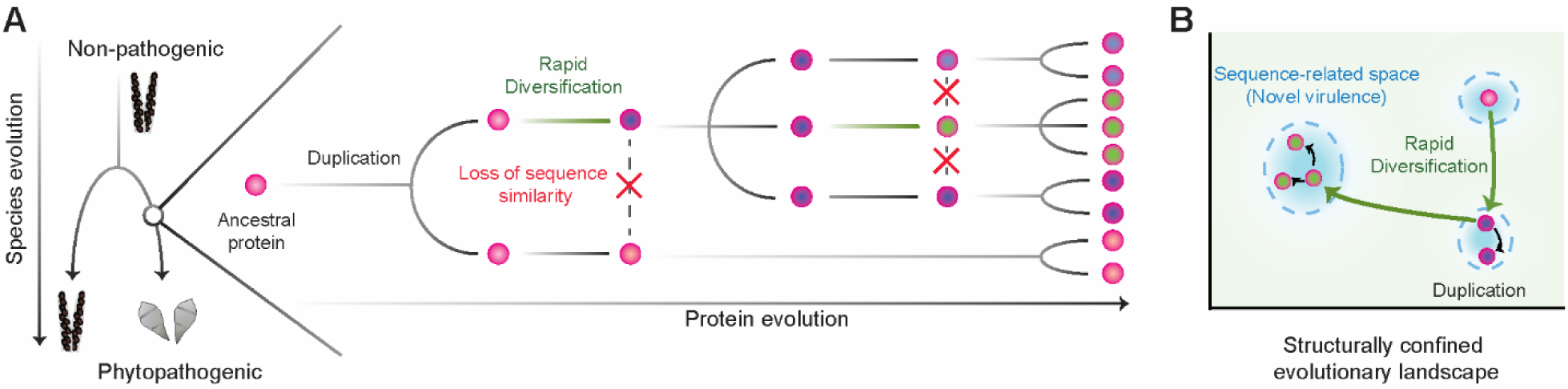
The divergent effector evolution. **A** The proposed evolution of a phytopathogen from an ancestral non-phytopathogenic species and effector families from ancestral proteins A protein that were present in the ancestral species undergoes a duplication event. A paralog rapidly diverges and losses sequence similarity to the other paralog. Such processes occur repeatedly, leading to contemporary protein groups that are not related by their sequences. The evolution of sequence-unrelated structurally similar effectors. The proteins exist in a structurally confined space. Through rapid diversification, sequence-unrelated structurally similar effector emerges, occupying a novel sequence-confined space that may have distinct functions.

Sequence-unrelated structurally similar effectors likely have distinct virulent functions and roles. For instance, different localizations of hydrophobin-like effectors in *Pgt* may reflect biochemical or biological specialization (Dagvadorj et al., 2017; Zhao et al., 2018; Andac et al., 2020). TIN2, TIN4 and TIN5 of *U. maydis* contribute at a varying degree to virulence and tumor formation (Brefort et al., 2014). MAX effectors display unique surface properties, potentially suggesting distinct host targets (de Guillen et al., 2015). In this sense, the emergence of sequence-unrelated structurally similar effectors could be a natural outcome of functional divergence. The rapid sequence diversification that diminishes sequence similarities between homologs may result from accelerating neo-functionalization. Alternatively, acquiring novel and strong virulent functions may be only accomplished by accessing other sequence-confined evolutionary realms within the structurally confined evolutionary landscape (Fig. 7C). Rapid sequence divergence may be, therefore, a necessary process for functional diversification. Many secreted proteins are rich in cysteine residues, and their disulfide bonds were suggested to increase the protein structure stability against host proteases and oxidizing environments (Kamoun, 2006; Stergiopoulos and de Wit, 2009; Saunders et al., 2012). We also suggest that the disulfide bonds may efficiently restrict the effector diversification process into the structurally confined evolutionary landscape, allowing effectors to rapidly diversify while maintaining the core structures.

The protein structure space is more confined than the protein sequence space (Koehl and Levitt, 2002). Therefore, some structurally similar proteins may be the outcome of convergent evolution. However, recurring examples of linking pathogen effectors with non-pathogenic homologs make divergent evolution more plausible to explain the roots of effectors. Under the divergent evolution hypothesis, any secreted protein may evolve virulence functions and form sequence-unrelated structurally similar effectors. Finding more classes of such protein families would not be surprising. This also could imply that there are numerous different solutions to evolve pathogenicity. This model, then, could explain similar compositions of secretomes between evolutionarily closely related species even after structure-based clustering; common presence of virulence factor-containing clusters shared between pathogens and non-pathogens; the existence of highly expanded effector groups, such as the RNAse and hydrophobin families in *Bgh* and *Pgt*, that originated from conserved secreted proteins; and most importantly, numerous, independent emergence of fungal parasitism. However, the divergent evolution model poses challenges in that a single, definitive role cannot be assigned to each effector family. Rigorous molecular biology, guided by structure-based evolutionary studies, will remain essential to deepen our understanding on functional divergence and unique utilization of effectors.

### Finer species sampling may reveal a closer evolutionary viewpoint for the existence of ancestral effector folds

We believe that ancestral origins of many nearly or entirely species-specific effector families can be revealed through finer sampling of fungal species. Although MAX effectors were nearly exclusive to *M. oryzae* in our study, *V. inaequalis* in the order Pleosporales of Ascomycota encodes many MAX effectors (Rocafort et al., 2022). Similarly, *Sporisorium reilianum* related to *U. maydis* encodes Tin2-like effectors found to be exclusive in *U. maydis* in our study (Tanaka et al., 2019). Even yeast-like *Pseudozyma hubeiensis* believed to be non-pathogenic seemed to share homologs (e.g., XP_012191528.1) (Konishi et al., 2013; Sharma et al., 2019). Therefore, resolving ancestral origins of all effector folds will require a larger scale comparative study.

### Singletons: missed prediction or true singletons

Although our study revealed interesting features of effector evolution, it may not yet provide a comprehensive perspective of pathogen effectoromes. Some putative virulence functions may have not been properly annotated or were filtered out based on the false prediction of signal peptides, leading to underestimation of effector family sizes. 7,207 (27%) secreted proteins remained as singletons and were therefore not discussed. The majority of these singletons are missing predicted structures. Some of these singletons may be true singletons without any evolutionarily related proteins in other species or within the species; Others may not be. For instance, a recent structural genomics study highlighted FOLD effectors in *F. oxysporum* f. sp. *lycopersici*, represented with Avr1 (SIX4), Avr3 (SIX1), SIX6 and SIX13 (Yu et al., 2021). Some of the proteins were not modeled with pTM scores > 0.5, and many of putative FOLD effectors remained as singletons in our study. This potentially suggests that some structural folds may be harder to predict with AlphaFold, and many potentially expanded, novel folds may be hidden in the singletons. Bechmarking AlphaFold’s prediction with experimentally determined structures could reveal that the folds of effectors could still be predicted with relatively low estimated precision (Yu et al., 2021). Experimentally determining protein structures of the singletons and including them in the training of AlphaFold may help to rescue some of the singletons in protein structure prediction.

### Disordered proteins: understudied new players of pathogenesis

Disordered fungal effectors have not been extensively studied. However, AVR-Rmg8 (109 amino acids) from *M. oryzae* (Anh et al., 2018) and Rsp3 (869 amino acids) from *U. maydis* (Ma et al., 2018) support the idea that disordered proteins have virulence roles and can be targeted by plant immunity. We found evidence that the core effector fold of *U. maydis* may be also diversifying by fusing with disordered stretches and subsequently contracting and extending them. This could be a strategy for a more rapid functional specialization than accumulating point mutations. At the same time, the intrinsically disordered regions may provide advantage to effectors, for instance, by aiding effector translocation (Marín et al., 2013). The additional importance may lie on the interaction between effectors and host immunity. The flexibility of the disordered regions and the absence of a single rigid conformation would not provide sufficient opportunities for the plant immune receptors to evolve specificity for. As sequence evolution occurs much faster on the long-disordered region (Brown et al., 2002), evading recognition could be accomplished more easily. Such features may drive the intrinsically disordered stretch to function as a shield of the core effector folds, reducing the frequency of the encounter between the core folds and immune receptors, while efficiently preventing recognition specificity development. Molecular biology will be an important avenue to elucidate how disordered effectors may function to compromise plant immunity.

### Predicted structures as a resource for future studies

The predicted structures generated in this study can serve as resources for larger comparative studies. The expansion to other fungal pathogens that infect humans, mammals and insects, as well as finely sampled non-pathogenic fungal species could illuminate distinct evolution of diverse lineages across the Fungal Kingdom. Structure-guided evolutionary study on plant-infecting bacteria, nematodes and insects may elucidate further insights on the plant-pathogen interactions.

## Methods

### Secretome prediction

The protein sequences of the species used in this study, excluding *M. oryzae* and *U. maydis*, were downloaded from the Joint Genome Institute (JGI) (Table S1) (Dean et al., 2005; Kämper et al., 2006; Espagne et al., 2008; Haas et al., 2009; Stajich et al., 2010; Amselem et al., 2011; Goodwin et al., 2011; Klosterman et al., 2011; Rouxel et al., 2011; de Wit et al., 2012; Manning et al., 2013; Wibberg et al., 2013; Grigoriev et al., 2014; Nemri et al., 2014; Baroncelli et al., 2018; Coradetti et al., 2018; DeIulio et al., 2018; Frantzeskakis et al., 2018; Kijpornyongpan et al., 2018; Li et al., 2019; Haridas et al., 2020). We used the neural network of SignalP v3.0 to identify secreted proteins (Dyrløv Bendtsen et al., 2004). The candidates were excluded if their predicted signal peptides overlapped with PFAM domains annotated with InterProscan v5.30-69.0 over 10 or more amino acids (Quevillon et al., 2005), or if their mature proteins contained any transmembrane helices detected with TMHMM v2.0 (Krogh et al., 2001). Only the mature proteins with 15 to 860 amino acids in length were selected for structure modeling.

### Structure prediction

The structures of 26,653 sequences were predicted by AlphaFold (Jumper et al., 2021). The full databases were used for multiple sequence alignment (MSA) construction with additional 1,689 fungal protein sequences downloaded from the JGI appended to the uniref90 database. All templates downloaded on July 20, 2021 were allowed for structural modeling. When the generated MSA was too large to process in our machine (>1G), we used HHfilter v3.3.0 to reduce the redundancy (Steinegger et al., 2019). For each protein, five models were generated with model_1, 3, 4 and 5, as well as model_2_ptm to obtain the pTM score. We selected the best model (ranked_0.pdb) determined by the average pLDDT score.

### Functional and structural annotations

The functional annotation was performed against Gene 3D v4.3.0, PFAM v33.1, Superfamily v1.75 with InterProscan v5.52-86.0 (Fox et al., 2014; Mistry et al., 2021; Sillitoe et al., 2019). We used Rupee for structural similarity search against SCOPe v2.07, CATH v4.3.0 and PDB chain databases downloaded on September 02, 2021 (TOP_ALIGNED, FULL_LENGTH) (Ayoub and Lee, 2019; Berman, 2000).

### Protein similarity searches

Sequence similarity searches were performed with BLASTP, HHblits and HHsearch for 26,653 secreted proteins (Camacho et al., 2009; Steinegger et al., 2019). HHblits and HHsearch require a profile. The profile was constructed by concatenating all MSAs produced by AlphaFold and filtering the concatenated MSA with HHfilter (-id 90 -cov 50 -maxseq 20000). All sequence similarity search outputs were filtered based on E-value ≤ 1E-10 and bidirectional coverage ≥ 65% prior to clustering. Structural similarity search was performed with TM-align (Zhang et al., 2005). Structural similarity was considered significant only if the pair of structures were predicted with pTM scores > 0.5, and their structural similarity were measured with TM-score > 0.5 normalized for both structures. The parameters were set more stringent than the criteria used in our previous work to reduce false clustering (Seong and Krasileva, 2021). To identify the RNAse supercluster, the previously used parameters were adopted for the *B. graminis* secretome: E-value < 10E-4 and bidirectional coverage > 50% for sequence similarity searches, and TM scores were > 0.5 for both structures or > 0.6 and > 0.4 for each structure for structural similarity searches.

### Clustering and network analysis

Protein clustering was performed sequentially with the similarity search outputs from BLASTP, HHblits, HHsearch and TM-align. Unlike our previous study that relied on a connected network (Seong and Krasileva, 2021), we applied the Markov clustering algorithm to reduce false clustering (Van Dongen, 2000). We first generated with mcxload a network of protein sequences based on -log_10_ (E-value) as weights from the BLASTP similarity search results, while capping the weight at 200 (--stream-mirror -- stream-neg-log10 -stream-tf ‘ceil(200)’). We then defined clusters with mcl with an inflation factor of 2 (-I 2.0). Once the protein sequences are assigned into clusters, the pairwise sequence similarity search outputs from HHblits were redefined to indicate the connectivity between the clusters. The average weights between the members in two clusters was used as the weight between the clusters, similarly to the average-linkage clustering. These processes were repeated for HHsearch and TM-align similarity search results as well. For structure-based clustering, as the TM-scores range from 0 to 1, these scores were used directly without converting them to a log scale.

### Motif analysis

The Y/F/WxC motif was identified by scanning 3-mer in a sliding window in the first 45 amino acids of the secreted proteins (Godfrey et al., 2010).

## Acknowledgements

We thank Dr. Brian Staskawicz for the computational resources, and Dr. Joseph W. Spatafora and the 1000 Fungal Genomes Project for allowing us to use the *Spathularia flavida* genome. We thank Chandler Sutherland, Dr. Daniil Prigozhin, Dr. Erin Baggs and Pierre Joubert for the critical review on the manuscript. This research relied on the Savio computational cluster resource provided by the Berkeley Research Computing program at the University of California. Kyungyong Seong is supported by the Berkeley Fellowship. Ksenia V Krasileva is supported by the Gordon and Betty Moore Foundation (grant number: 8802) as well as by the joint funding from the Foundation for Food and Agriculture and 2Blades (CA19-SS-0000000046) and the Innovative Genomics Institute.

## Author Contributions

K.S. conceived and conducted the research and wrote the manuscript. K.V.K. supervised the research.

## Data Availability

All the datasets and scripts used in this study can be downloaded from Zenodo: 10.5281/zenodo.6480453.

