## Supplementary material for "Comparative computational structural genomics highlights divergent evolution of fungal effectors": Figure S

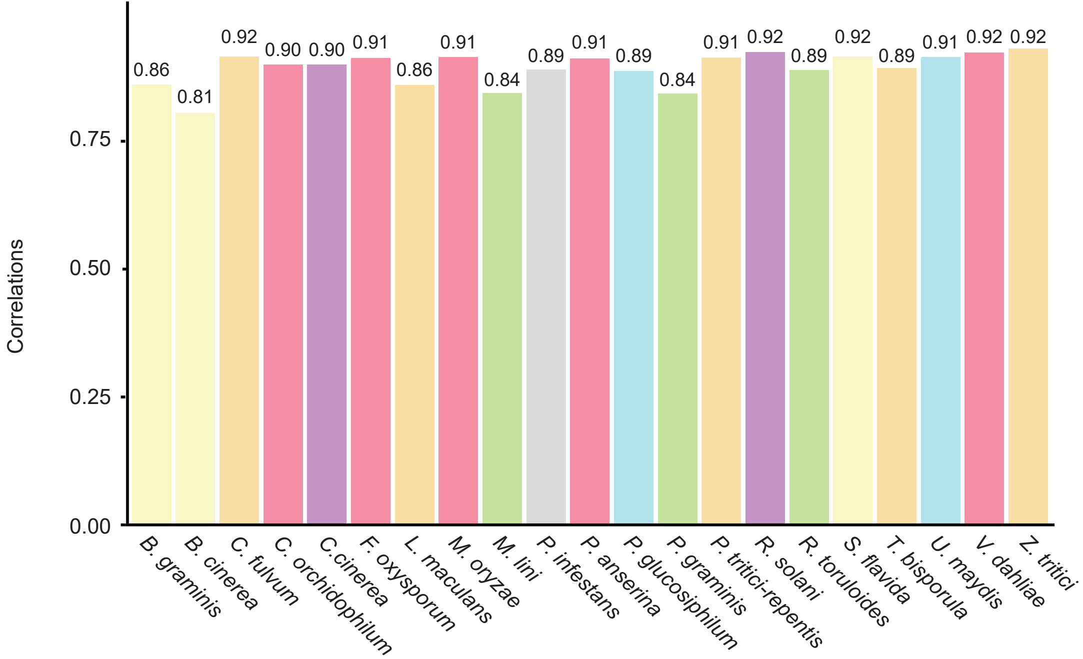


**Figure S1. The correlation between the global measure and average local measure of prediction accuracy.**

The global measure of prediction accuracy (pTM) was compared to the average local measure of prediction accuracy (pLDDT) for each structure. A Pearson correlation coefficient was calculated for each species.

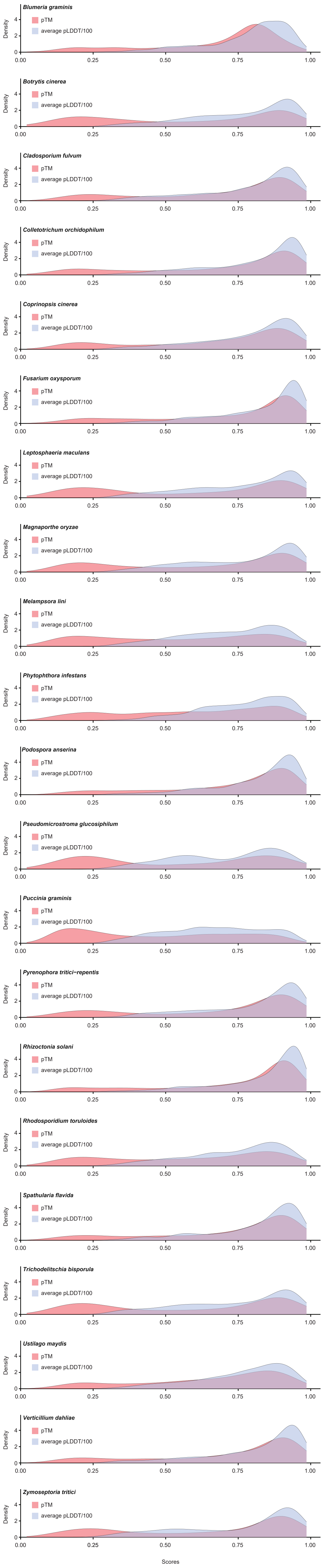


**Figure S2. The distribution of the global measure and average local measure of prediction accuracy.**

The distributions of the global measure of prediction accuracy (pTM) and the average local measure of prediction accuracy (pLDDT) are provided for each species. As the pLDDT ranges from 0 to 100, the pLDDT scores were divided by 100 to plot with the pTM scores.

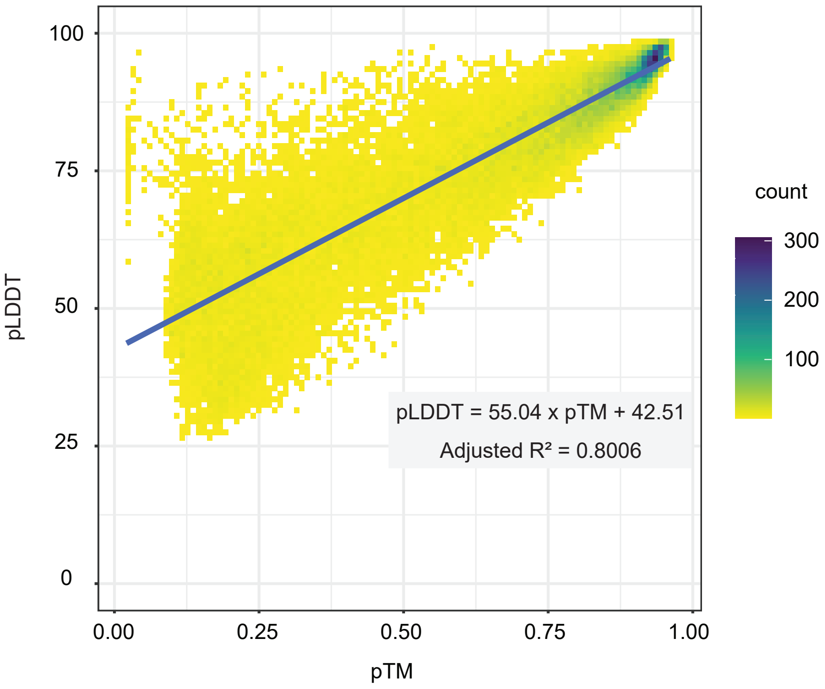


**Figure S3. The linear regression for pTM and pLDDT.**

A linear fit was found for the global measure of prediction accuracy (pTM) and the average local measure of prediction accuracy (pLDDT). Both coefficients were significant with p-value < 2 x $10^{-16}$.

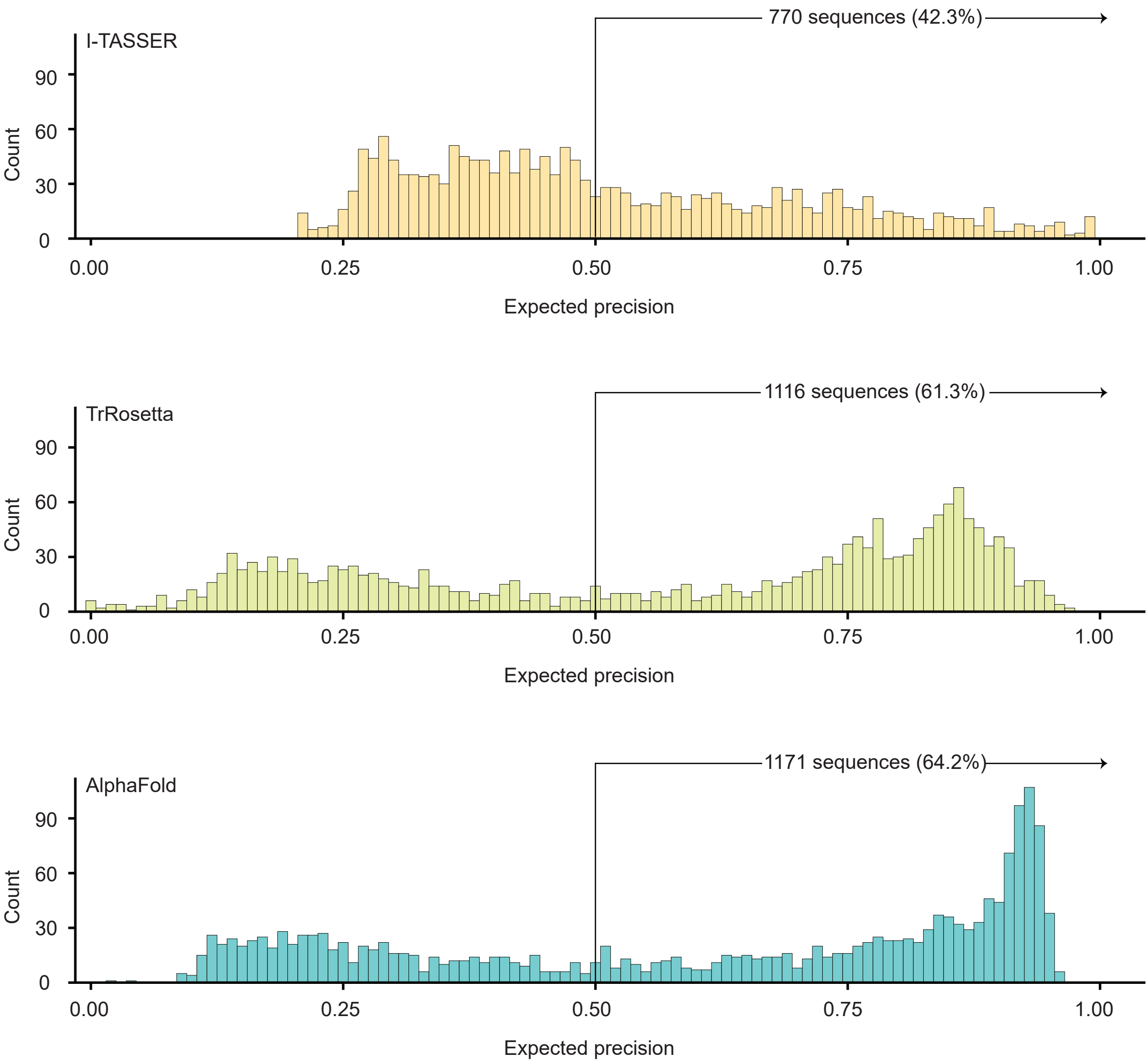


**Figure S4. The comparison of estimated precision from AlphaFold, TrRosetta and I-TASSER for the predicted structures of *Magnaporthe oryzae* proteins.**

The distribution of expected precision from the three structure prediction tools for the same set of 1,822 secreted proteins from *Magnaporthe oryzae*. The number of sequences, the structures of which were predicted with expected precision > 0.5, are indicated.

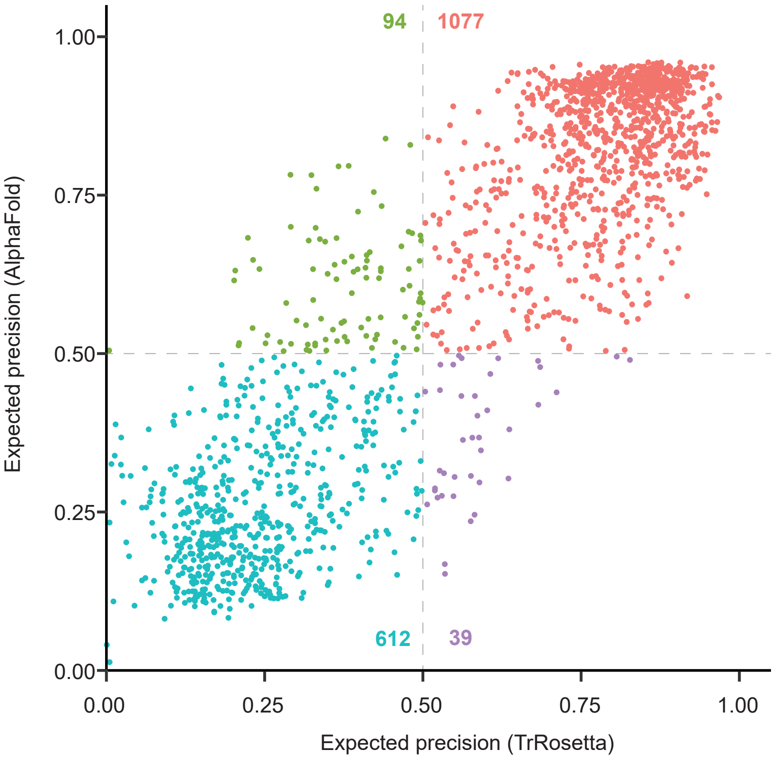


**Figure S5. The comparison of expected precision for AlphaFold and TrRosetta structures predicted for *Magnaporthe oryzae*'s secreted proteins.**

The distribution of expected precision from AlphaFold and TrRosetta for the same set of 1,822 secreted proteins from *Magnaporthe oryzae* were displayed in the Cartesian coordinate. The number of sequences that belong to each coordinate is indicated.

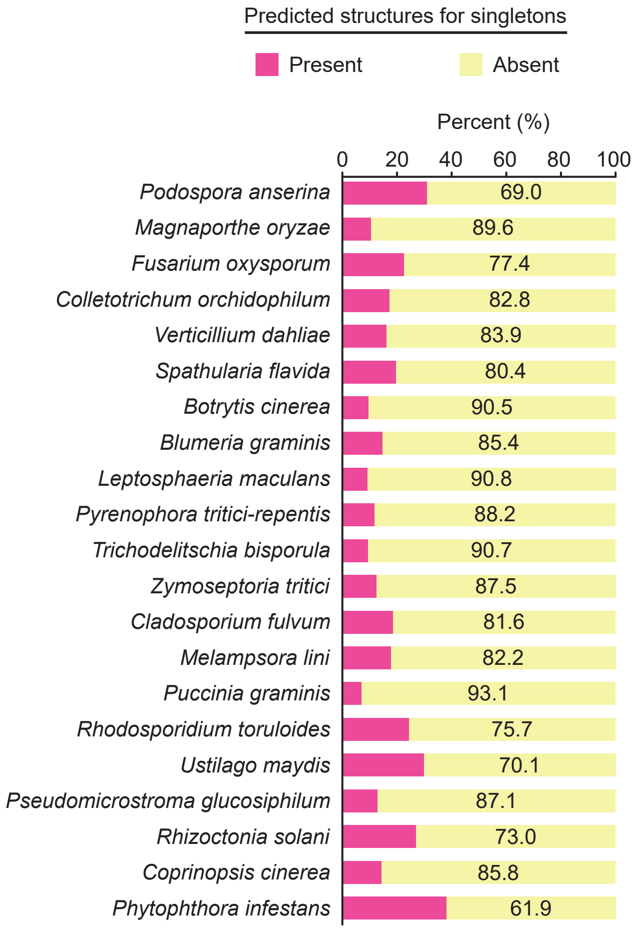


**Figure S6. The structure prediction statistics for singletons.**

The proportions of singletons in each species that have or do not have predicted structures are indicated. These singletons did not have any sequence or structure-related proteins in both individual and whole-secretome clustering.

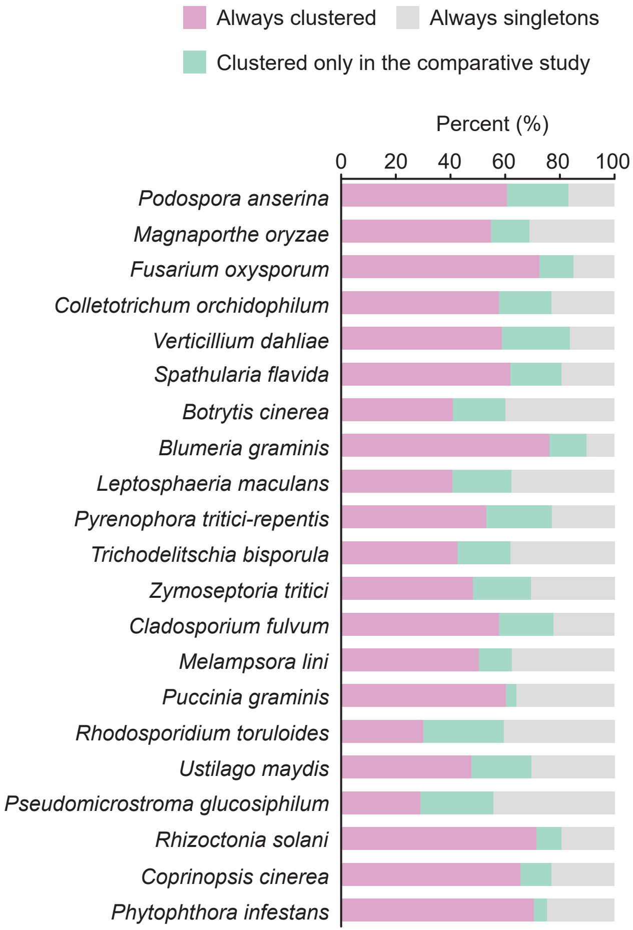


**Figure S7. The statistics of secretome clustering.**

The secretome of individual species or entire 21 species in this study were clustered based on sequence and structural similarity. The secreted proteins were categorized into 'Always clustered' if the proteins appear in a cluster in both individual and whole-secretome clustering outputs, 'Clustered only in the comparative study' if the proteins appear in a cluster only in the whole-secretome clustering output, and 'Always singletons' if the proteins appear as singletons.

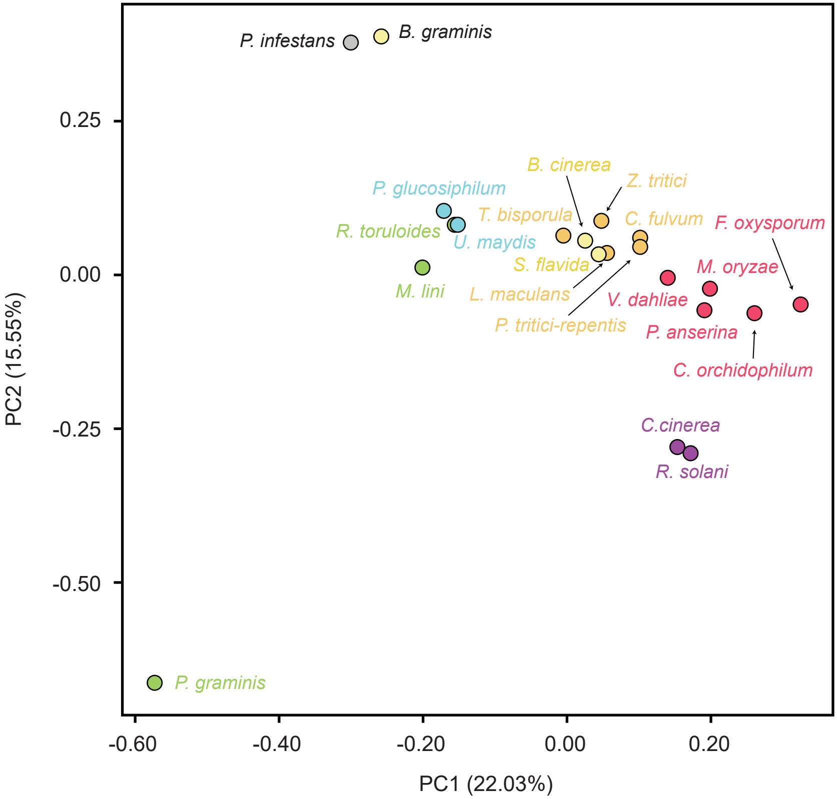


**Figure S8. The principal component analysis on BLAST-based clustering outputs.**

The secretome of the 21 species was clustered based on sequence similarity identified with BLASTP. The scaled centered cluster counts were reduced into the two dimension with the principal component analysis. The color of the circles is given based on the taxonomy (class or subphylum) following Figure 1A.

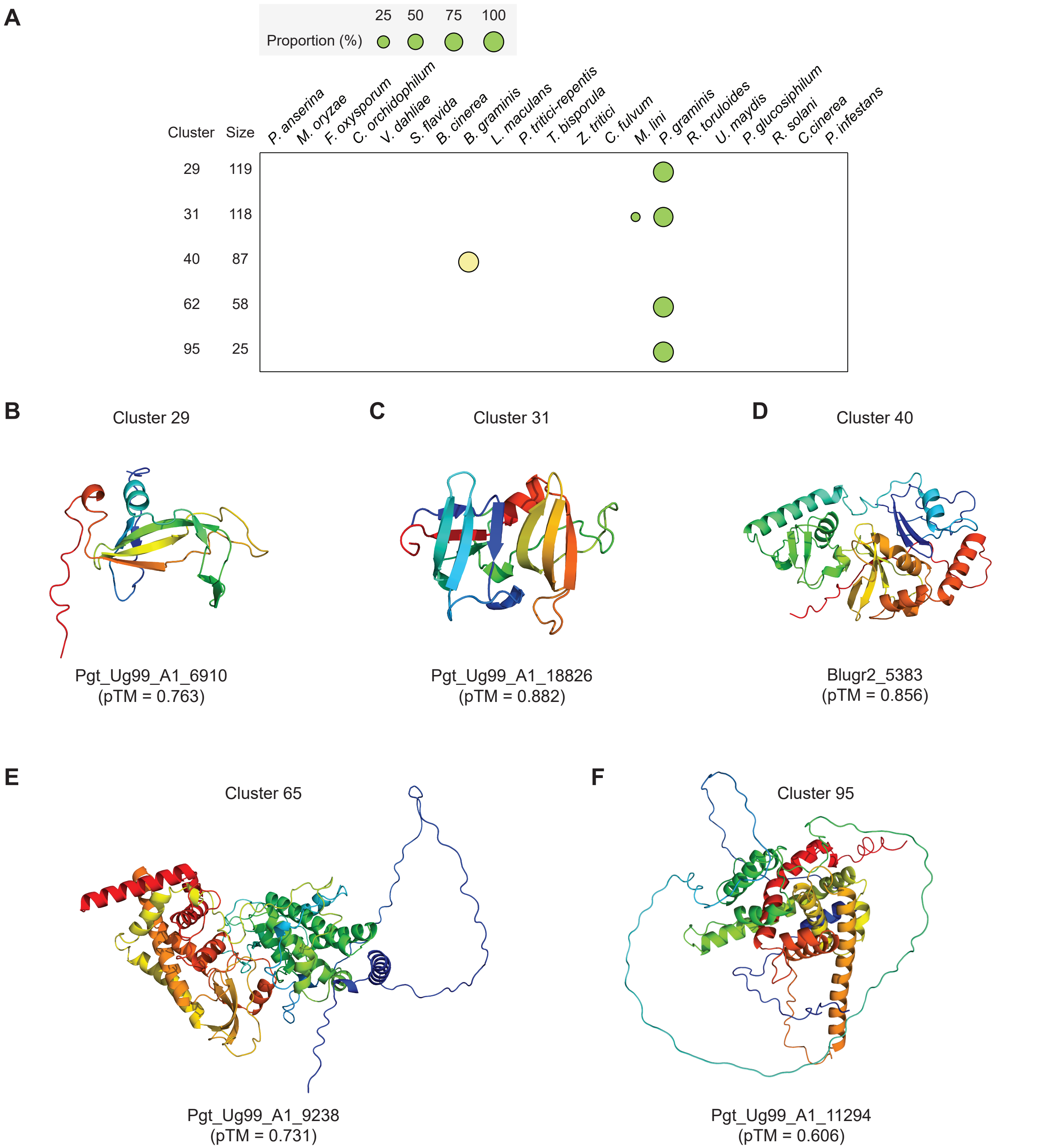


**Figure S9. Nearly or entirely species-specific clusters without known virulence factors.**

**A**. Nearly or entirely species-specific clusters found in *Blumeria graminis* and *Puccinia graminis*. The relative composition of each species is indicated with circles with varying sizes. **B** – **F**. The selected predicted structures from each cluster.

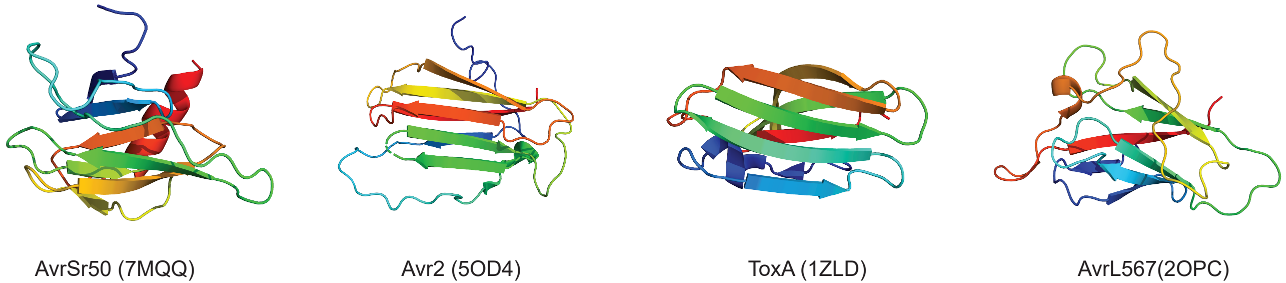


**Figure S10. Selected effectors that contain β-sandwich-like folds.**

Experimentally determined structures of AvrSr50 (Cluster 171), Avr2 (Cluster 68), ToxA (Cluster 68) and AvrL567 (Cluster 1283). The PDB accessions are given in the parentheses.

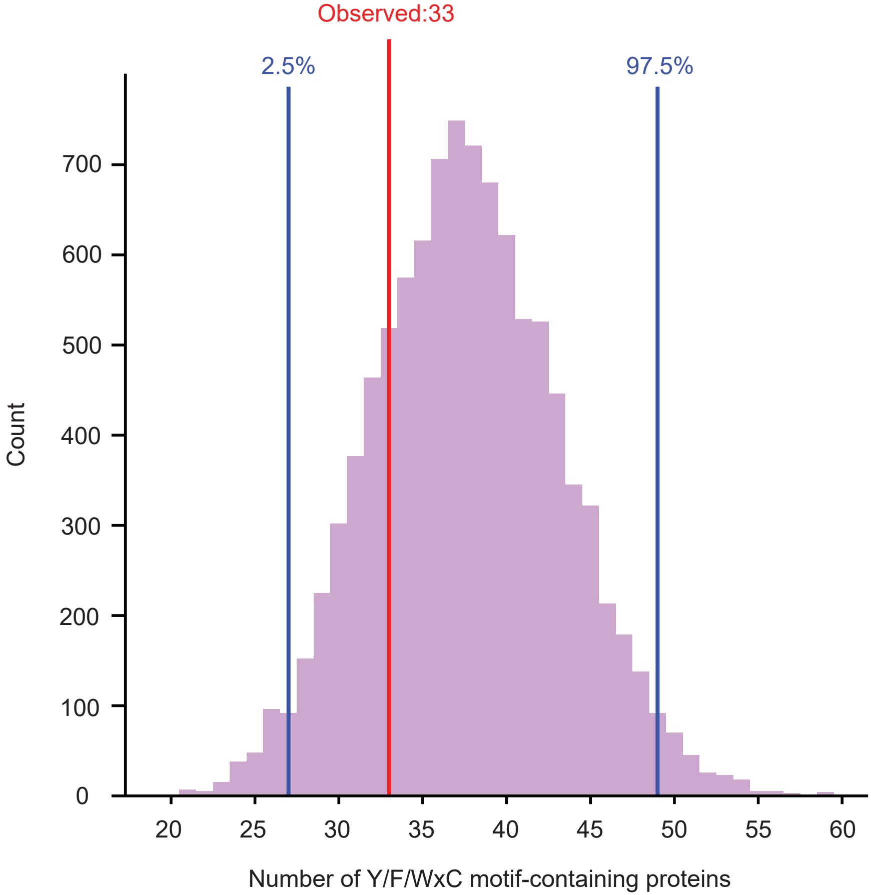


**Figure S11. The distribution of the number of Y/F/WxC motifs in the permutation test for non-RNAses in *Blumeria graminis*.**

For secreted proteins from Blumeria graminis that do not belong to the RNAse supercluster, the first 45 amino acids were permutated and the Y/F/WxC motif was searched. This permutation test was performed 10,000 times, and the distribution of the number of Y/F/WxC motif-containing sequences is provided. The observed 33 Y/F/WxC motif-containing sequences could occur by chance.

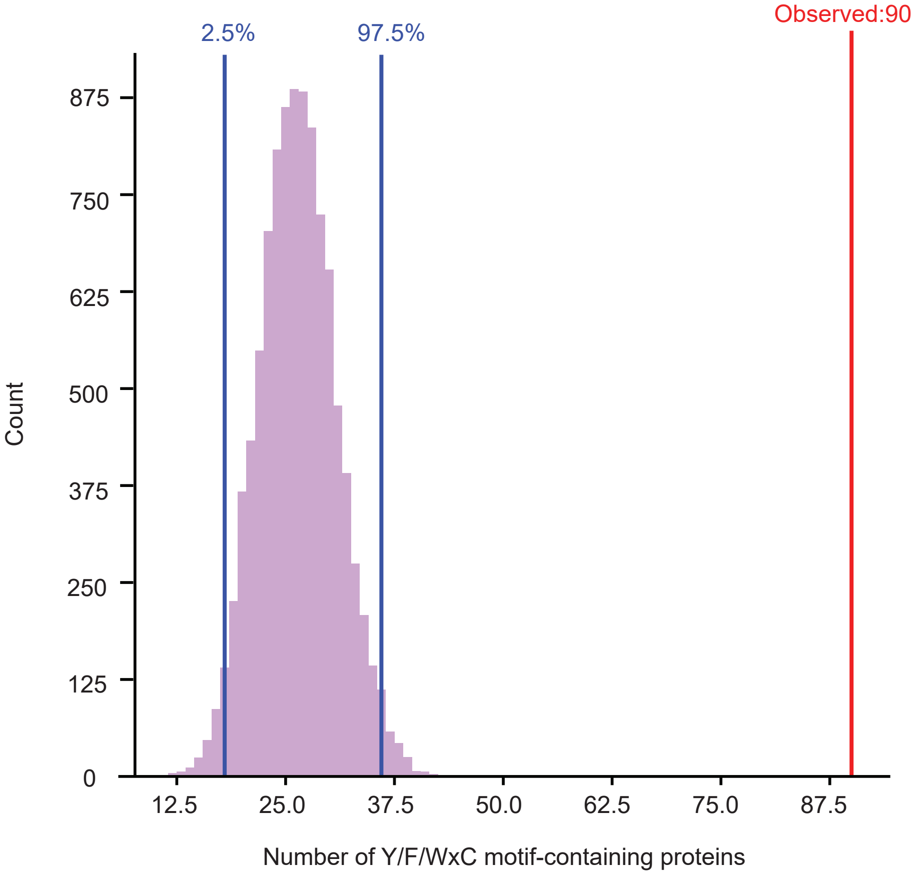


**Figure S12. The distribution of the number of Y/F/WxC motifs in the permutation test for Cluster 25 members in *Puccinia graminis*.**

For Cluster 25 members from *Puccinia graminis*, the first 45 amino acids were permutated and the Y/F/WxC motif was searched. This permutation test was performed 10,000 times, and the distribution of the number of Y/F/WxC motif-containing sequences is provided. The observed 90 Y/F/WxC motif-containing sequences unlikely occur by chance.

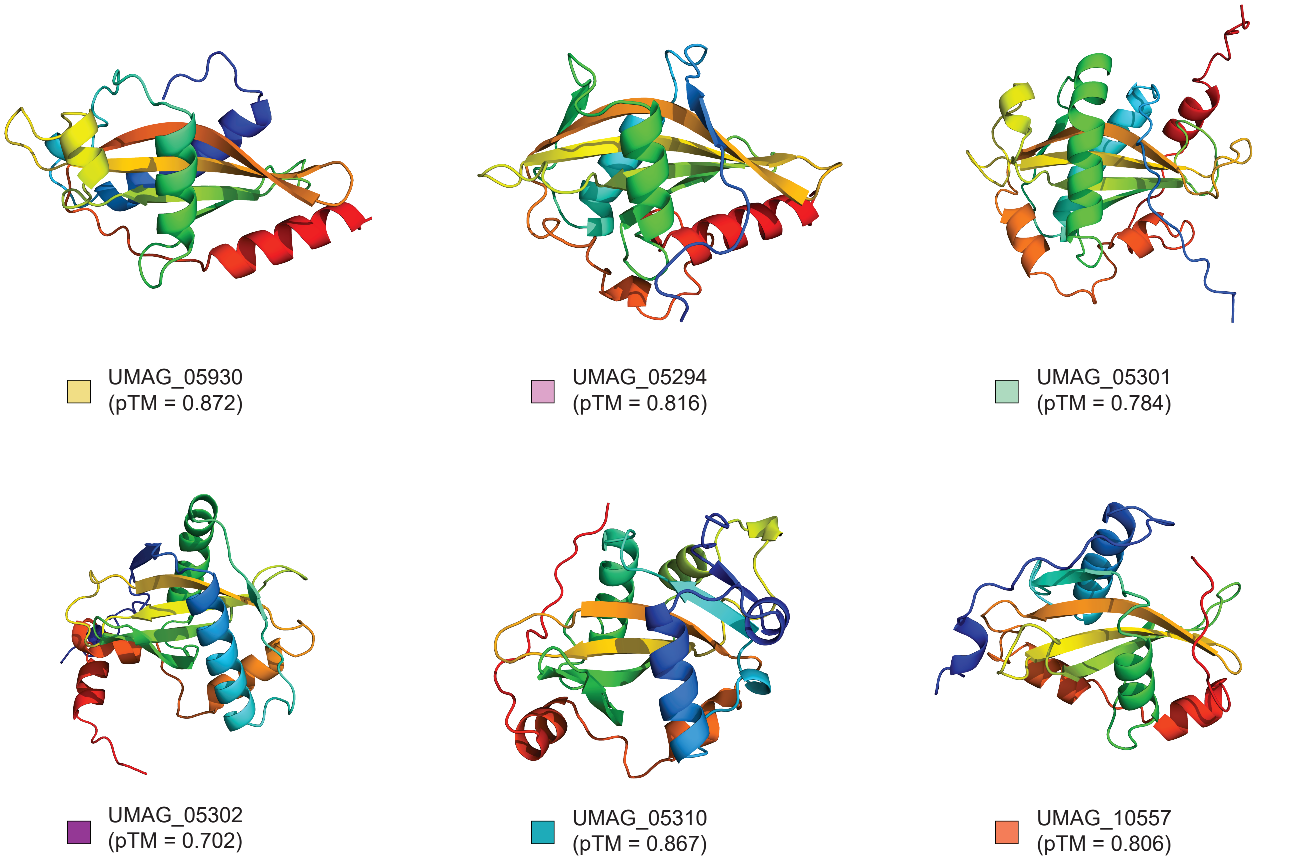


**Figure S13. The selected structures of the Tin2-like effectors.**

A representative structure was selected from each subcluster of Cluster 96. The membership of the sequences is indicated with colored boxes. The colors are the same as in Figure 5.

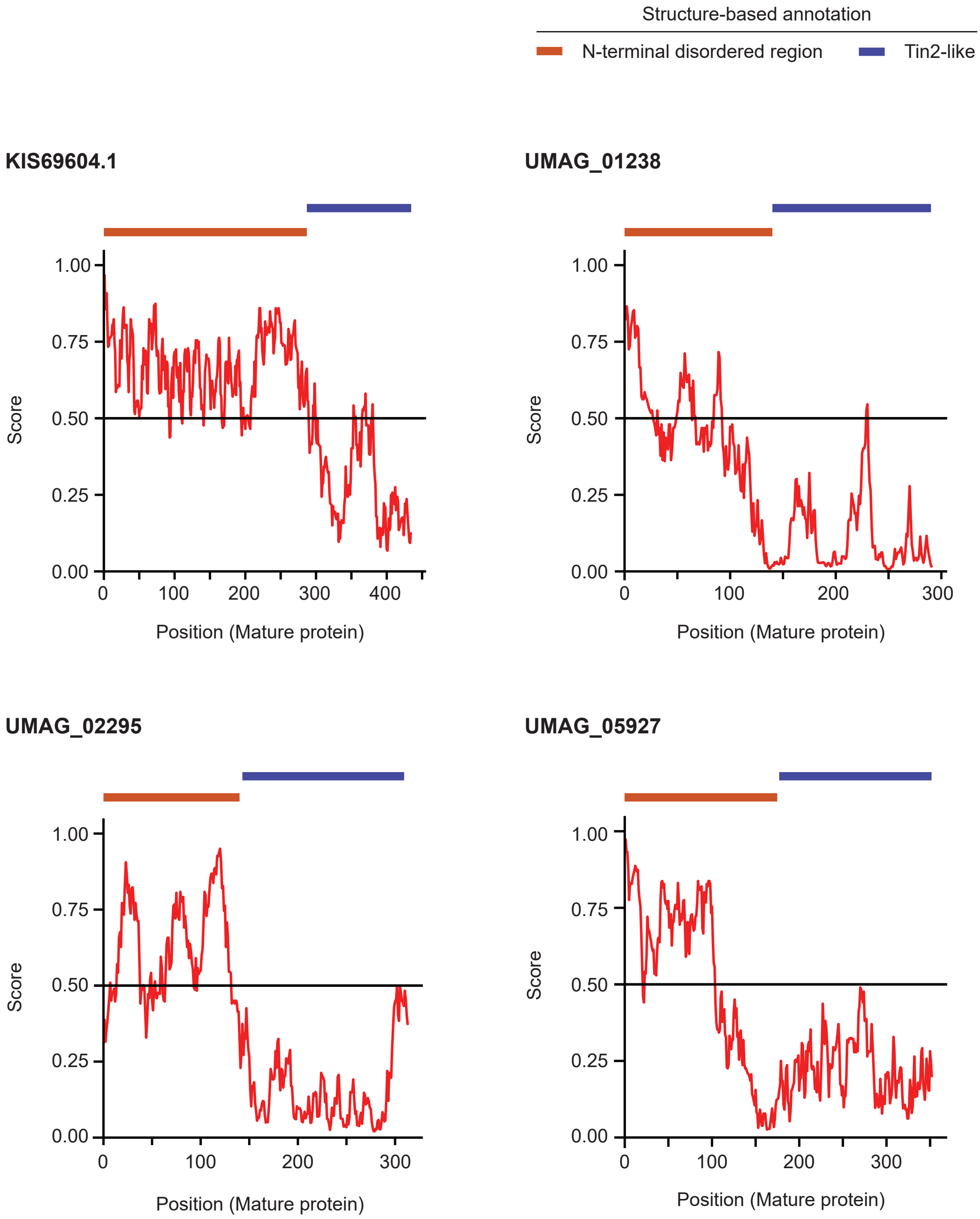


**Figure S14. The prediction of the disordered regions for the Tin2 fusion proteins.**

IUPred2A was used to predict the intrinsically disordered regions for the Tin2 fusion proteins given in Figure 5. The structure-based disordered region and Tin2 fold are indicated with orange and blue bars on top of the plot. The score > 0.50 was used to determine the intrinsically disordered regions.

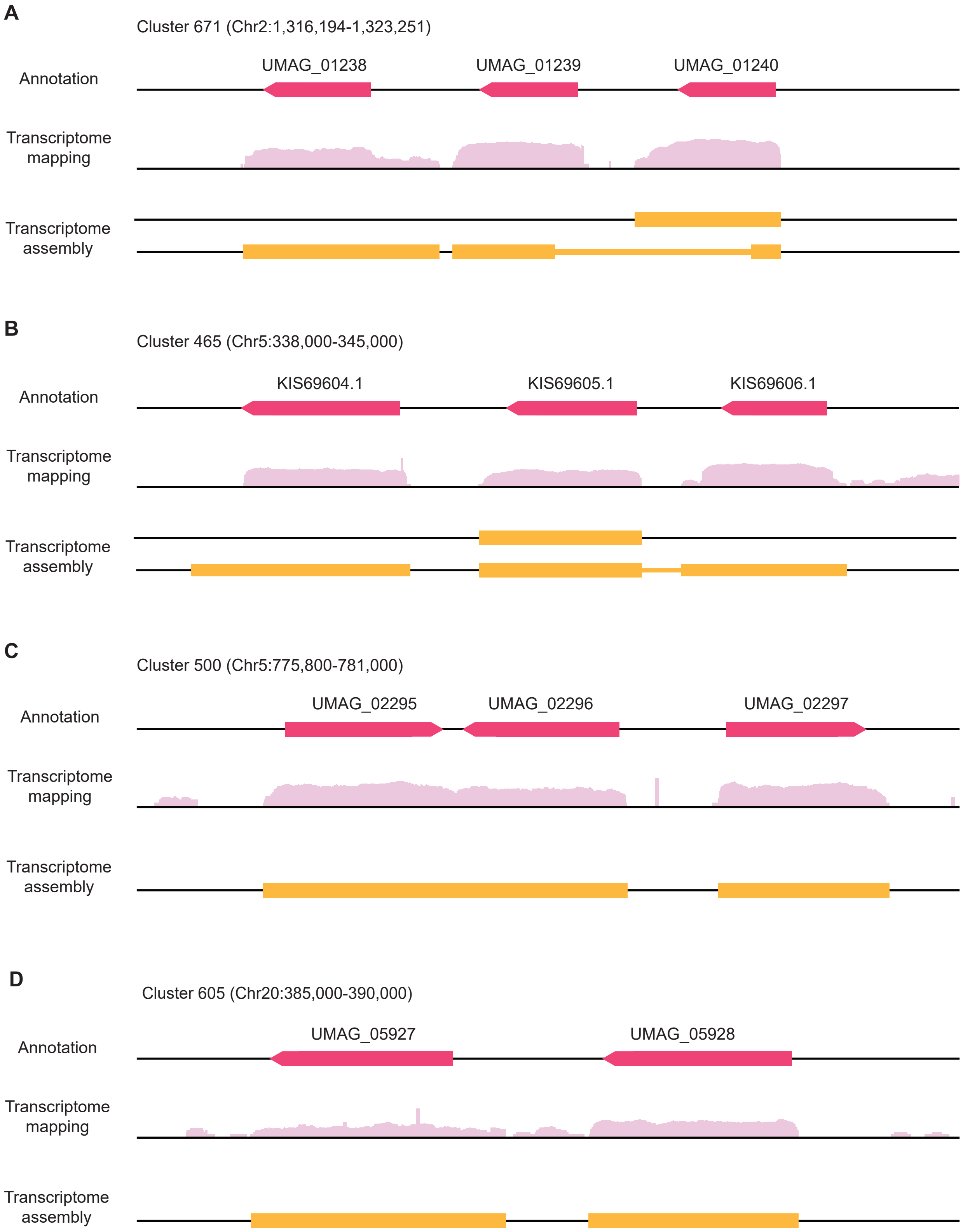


**Figure S15. Transcriptomic mapping and assembly results for the Tin2 fusion proteins.**

A subset of publicly available transcriptomic data was aligned or assembled to examine the annotation quality of the Tin2 fusion proteins. The results are displayed for some Tin2 fusion proteins that exist in proximity.

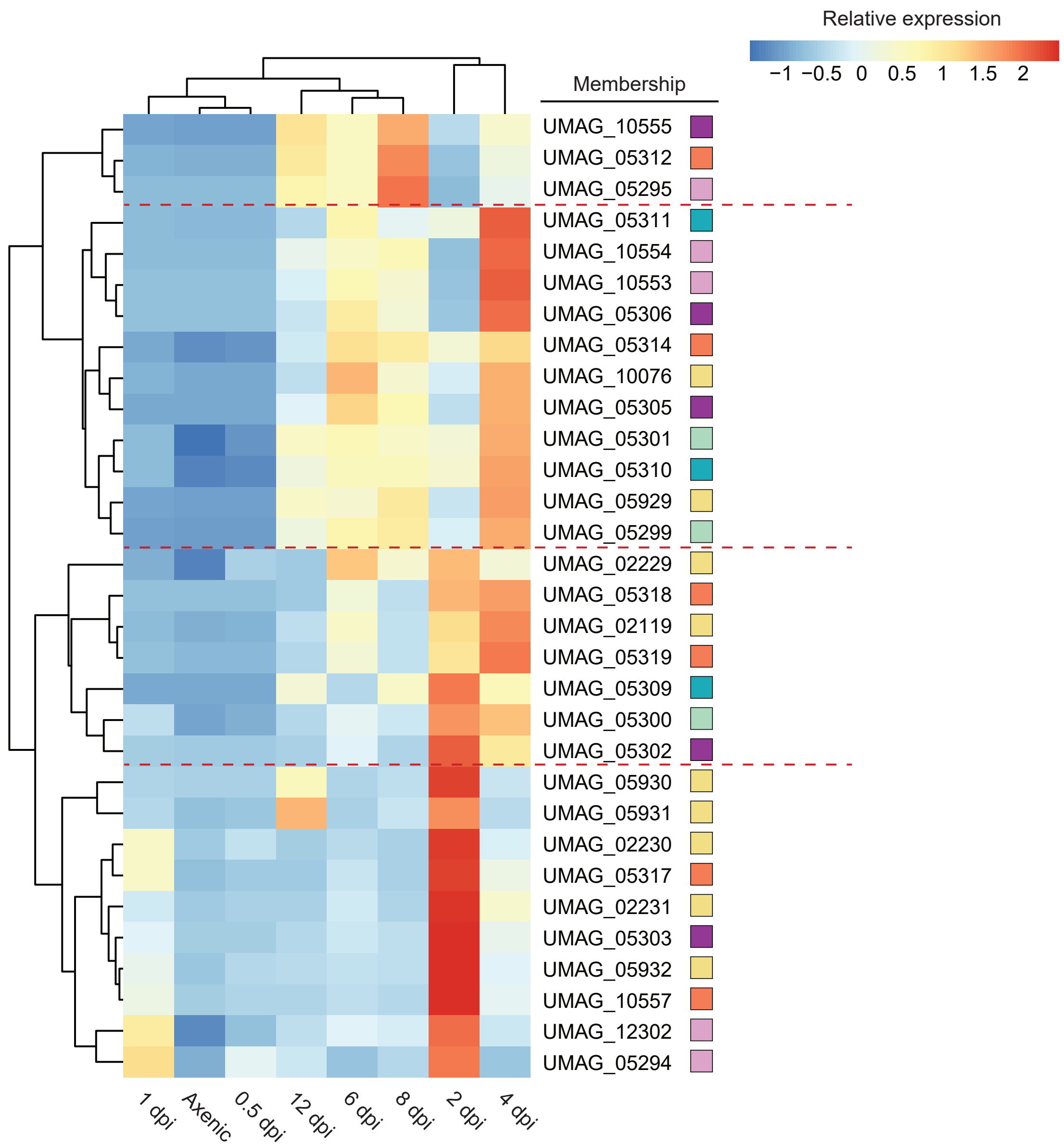


**Figure S16. Hierarchical clustering of the core Tin2-like cluster (Cluster 96).**

The expression profiles of the Cluster 96 members were clustered with hierachical clustering. The membership of each sequence is indicated with colored boxes. The colors are the same as in Figure 5.
